# Lithocholic acid modulates the growth of butyrate-producing bacteria and is decreased in the feces of stunted children

**DOI:** 10.64898/2026.05.04.722639

**Authors:** Kelsey E. Huus, Julian R. Garneau, Nermin Akduman, Simon Yersin, Jun Han, Mariia A. Beliaeva, Cordula Gekeler, Leonardo Boldt, Munir Winkel, Christoph H. Borchers, B. Brett Finlay, Michael Zimmermann, Philippe J. Sansonetti, Lisa Maier, Pascale Vonaesch, the Afribiota Investigators

## Abstract

Bile acids modulate the intestinal microbiota and serve as key signaling molecules in host physiology. Bile acid dysregulation has been implicated in nutritional and inflammatory diseases; however, data on the pool of bile acids present in stunted children or children suffering of environmental enteric dysfunction (EED) is limited, particularly in the upper intestinal compartment where disease phenotypes are most relevant. In this study, we performed a targeted metabolomics approach on 75 bile acids and their derivatives, including gastric and duodenal aspirates and fecal samples from almost 1000 children from two Sub-Saharan cities. We found that levels of secondary bile acids, especially lithocholic acid, are significantly lower in the feces of stunted and EED children, while ursocholic acid and its derivatives are significantly higher. Levels of primary and sulfated bile acids are also increased in the feces of children with EED. Microbiota sequencing revealed that high lithocholic acid levels are positively associated with butyrate-producing bacteria, while negatively associated with oral taxa like *Streptococcus* and *Veillonella*. *In vitro* tests on a panel of reference strains showed that oral bacteria bioaccumulate and are inhibited by a variety of bile acids, while lithocholic and chenodeoxycholic acids modulate the growth of several butyrate-producing bacteria. This effect was even stronger with tauro- or glycol-conjugated bile acids. Exposing stool-derived *in vitro* communities from children in Afribiota to these bile acids confirmed their positive impact on butyrate producers and negative effect on overgrowing oral taxa. Our findings suggest that secondary bile acids, reduced in stunting and EED, modulate the growth of butyrate-producing bacteria while suppressing harmful oral taxa, highlighting their potential as tools to modulate microbiota composition.

## Introduction

Bile acids are key molecules in human gastrointestinal health. They are implicated in protein digestion and lipid absorption and in cellular signaling through the farnesoid X receptor (FXR), the G-coupled protein receptor 5 (TGR5), the c-Jun amino terminal kinase (JNK) pathway as well in several other pathways; these pathways regulate in their turn genes for body temperature, appetite and overall energy homeostasis (reviewed in^1–4^). Bile acids are thus key metabolites in nutrition-associated diseases, including obesity and metabolic disease^2,4–8^, and have also been implicated in child undernutrition^9,10^.

Bile acids and their derivatives are also known to shape the microbiome, as some bile acids are toxic to certain members of the microbial community^11,12^. Indeed, biliary obstruction is associated with bacterial overgrowth and translocation of bacteria in the small intestinal tract^13^, and administration of bile acids has been shown to alleviate this phenomenon^14^. Bile acids are also directly involved in regulating the virulence of enteropathogens^12,15–18^, in maintaining colonization resistance against invading bacteria^19–22^, in inflammation^1,23^, peristalsis and gut permeability^13,14^ within the gastrointestinal tract.

Bile acids are mainly produced by the human liver by oxidation of cholesterol and are then conjugated with either glycine or taurine (reviewed in^1,4^). The main primary bile acids produced in the liver are the glyco- and tauro-conjugates of cholic, chenodeoxycholic, and deoxycholic acids. Once secreted in the intestinal tract, they are then readily modified into many different secondary bile acid by the human microbiome. Typical modifications include deconjugation, 7-α-dehydroxylation, oxidation, epimerization, amidation, esterification, sulfation, glucuronidation, and dehydration, to name just a few. The pool of bile acids is dictated by sex, age, the intestinal compartment, diurnal rhythm, and overall health status. The unconjugated bile acids are reabsorbed before they reach the terminal ileum and circulate back to the liver via the enterohepatic system (reviewed in^1,4^).

The pool of bile acids is affected in many diseases. In inflammatory bowel disease (IBD), there is an increase of *Enterobacteriaceae* and a decrease in *Bacillota* (formarly called *Firmicutes*), including *Clostridiales*, in the feces, which is associated with increased luminal levels of primary bile acids, higher levels of sulfated bile acids, and a decrease in luminal secondary bile acids^23–26^. Furthermore, the secondary bile acid lithocholic acid is key in providing colonization resistance against *C. difficile*^19–22^, and changes in the pool of bile acids are thus also often accompanied by overgrowth of given (pathogenic) bacteria.

Childhood stunting is a syndrome affecting roughly one-fourth of all children below the age of five years across the globe^27^ and is a syndrome that is associated with changes in the intestinal microbiome composition^28–33^. The microbial disease signature is characterized through three main hallmarks: overgrowth by bacteria of oral origin in the small intestinal space (SIOBO)^29–32^, overrepresentation of enteropathogens, especially *Campylobacter* spp.^28,31,32^ and *Shigella* spp.^34^ and a reduction of butyrate-producing *Clostridiales* in the colon^28,32^. Stunting is closely associated with an inflammatory disease of the small intestine, called environmental enteric dysfunction (EED). EED is characterized by a reduction in the absorptive villi of the small intestinal tract, increased gut permeability as well as influx of immune cells^35^. While non-invasive biomarkers of EED have been shown to correlate inconsistently with one another and with stunting^36–38^, there seems to be consensus that markers of intestinal inflammation, including α-1-antitrypsin and calprotectin are associated with subsequent linear growth^38^.

Prior studies have shown changes in the pool of bile acids in the context of childhood undernutrition and EED, focusing mainly on serum levels^39–41^. In these studies, the fecal bile acid pool and its association with given microbial signatures was not studied.

Gram-positive bacteria, including many oral species associated with stunting, are known to be more susceptible to bile acids compared to gram-negative bacteria^43^. Furthermore, many members of the *Clostridiales* that are reduced in stunted children are implicated in 7-α-dehydroxylation of bile acids (reviewed in^12^). We thus hypothesized that stunted child growth and EED might be associated with changes in the pool of intestinal bile acids, both leading to and sustaining microbial dysbiosis and overall pathophysiology within the host.

To explore this hypothesis, we assessed the pool of bile acids in the stomach, duodenum and feces of over 900 children living in two of the most affected regions of Sub-Saharan Africa and correlated the relative abundances of the bile acids with clinical factors, including nutrition and inflammation as well as with the composition of the microbiota. We then explored the effect of the bile acids on the microbiome composition *in vitro* using large-scale screening and metabolomics analysis on a core set of reference strains of the human fecal microbiome as well as on stool-derived communities from children from Sub-Saharan Africa.

Our results show key changes in the bile acid pool of both stunted children and children with EED. Furthermore, we show that the changes in these bile acids correlate with EED-associated taxa in the human cohort. Experimentally, we show that the addition of these key bile acids leads to inhibition of oral bacteria and boosted growth of *Lachnospiraceae* both in single cultures as well as in complex communities, thus partially explaining the observed dysbiosis in stunted child growth and EED.

## Material and methods

### Study group

The children and samples included in this study are part of the Afribiota project, a cross-sectional study on stunting performed in children aged 2-5 years in Antananarivo, Madagascar, and Bangui, Central African Republic^42^. All samples from children with available bile acid data and duodenal samples with a pH of >5 and gastric samples with a pH <4 were considered for the final analysis. When applicable, data available for microbiota profiling were included in the final analysis. The samples included are summarized in the flowchart in Supplementary Figure S1. The detailed study protocol^42^ as well as the associated metadata on social and environmental factors^43^, biomarkers^36^, enteropathogen carriage^30,44^, parasite carriage^45^, and microbiota composition^30–32,46,47^ have been reported previously. Briefly, children living in specific districts of Bangui, Central African Republic or Antananarivo, Madagascar, not experiencing any severe disease such as acute respiratory distress, HIV, watery or bloody diarrhea or acute undernutrition and aged 2-5 years were included. In each country, the target size was 460 children, classified as severely stunted, moderately stunted and not stunted according to the median height of the WHO reference population^48^. Metadata collected include nutritional status, age, socio-economic factors, iron and hemoglobin levels, biomarkers of intestinal permeability and mucosal and systemic inflammation as well as parasite and enteropathogen load. Metadata was collected using a standardized questionnaire and routine diagnostics as previously described^36,43^. We collected feces for all subjects. For undernourished subjects, we also collected duodenal and gastric aspirates. Caregivers were instructed to collect the feces in the morning before coming to the inclusion visit. Gastric and duodenal samples were collected using a paediatric nasogastric tube (Vygon, France) and were only collected for stunted children (ethical constraint). Once the gastrointestinal samples were collected, they were aliquoted, frozen at -20 °C, and transferred the same day to a - 80 °C freezer (Bangui) or directly snap frozen in liquid nitrogen and then transferred to a -80 °C freezer within a week (Antananarivo). Biobanking was performed by the Clinical Investigation and Access to BioResources Platform (ICAReB) at the Pasteur Institute, Paris, and by the Pasteur Institutes of Madagascar and Bangui. The study protocol for AFRIBIOTA has been approved by the Institutional Review Board of the Institut Pasteur (2016-06/IRB) as well as the National Ethical Review Boards of Madagascar (55/MSANP/CE, May 19^th^, 2015) and CAR (173/UB/FACSS/CSCVPER/16). Informed consent was obtained from all subjects involved in the study.

### Bile acid profiling

To measure bile acid content, 100 mg of frozen material of each gastric, duodenal, or fecal sample was aliquoted in an Eppendorf tube and subsequently lyophilized. All bile acids were reported to dry mass weight. Bile acids were measured at the Genome British Columbia Proteomics Center, the University of Victoria, BC, Canada. The lyophilizates were dissolved in 70% aqueous acetonitrile at a concentration of 20 μL per mg of dry weight. The samples were vortex-mixed for 30 s, sonicated in an ice-water bath for 5 min, and then centrifuged at 21,000 g and 10 °C for 10 min in an Eppendorf 5425 R centrifuge. For the fecal samples, 10 μL of the resultant clear supernatant from each sample was mixed with 50 μL of an isotope-labeled internal standard solution that contained 14-deuteraed bile acids as internal standards, and 0.94 mL of water inside polymeric reversed-phase Strata-X cartridges (60mg/1mL).

For the duodenal and gastric samples, 50 μL of the clear supernatant from each sample was mixed with 50 μL of the IS solution and 0.90 mL of water inside the cartridges. The reversed-phase solid-phase extraction cartridges were activated with 1-mL methanol and reconditioned with 1-mL water before sample loading. Under positive pressure, the flow-through solutions were discarded. The cartridges were washed with 1 mL of water twice. Bile acids were subsequently eluted with 1 mL of methanol-acetonitrile (1:1) twice under positive pressure. The collected and pooled fraction for each sample was dried down in a speed-vacuum concentrator. The dried residues were reconstituted in 100 μL of 40% methanol. Alongside the sample preparation, ten serially diluted calibration solutions containing standard substances of all the measured bile acids were prepared in the same IS solutions. The concentrations of each bile acid in the calibration solutions ranged from 0.000002 to 10 µM. 20 μL aliquots of the sample and calibration solutions were injected for the analysis on an LC-MS/MS system composed of an Agilent 1290 Infinity II UHPLC coupled to a Sciex 4000 QTRAP mass spectrometer. The MS instrument was operated in the multiple-reaction monitoring (MRM) mode with negative-ion (-) detection. A Waters BEH 1C18 UPLC column (2.1 mm I.D., 150-cm long and 1.7 μm particle size) was used for the chromatographic separation with the mobile phase composed of (A) 0.01% formic acid in water and (B) 0.01% formic acid in acetonitrile for binary-solvent gradient elution. The detailed description of the equipment and operating parameters is given in^51^, with necessary modifications. All the raw data files were recorded using Sciex Analyst® 1.7.3 and processed using the Sciex MultiQuant 1.2 software. Linear regression calibration curves of individual bile acids were constructed between analyte-to-IS peak area ratios (As/Ai) versus molar concentrations (nmol/mLµM) and were used for concentration calculation using the data acquired from the sample solutions for. The bile acids were treated over three consecutive batches, for which we controlled in the final data analysis. Names in between the three analysis rounds were harmonized and data merged accordingly. The total of bile acids in each sample was calculated as the sum of all the measured bile acids. Bile acids were analyzed individually and grouped as follows: conjugated bile acids were defined as the sum of all taurine or glycine-conjugated bile acids and their derivatives, non-conjugated bile acids as all other bile acids measured, bile acid sulfates (sulfo) as all bile acids that are sulfated, urso-derivatives as all bile acids with 7 alpha-OH epimerization and glucuronide bile acids (gluco) were defined as all bile acids with a glucuronide. The chenodeoxycholic acid (CDCA) pathway members include chenodeoxycholic acid, 3-oxo-chenodeoxycholic acid, 3-oxo-lithocholic acid, lithocholic acid, 7-oxo-chenodeoxycholic acid, and 7-oxolithocholic acid. Members of the cholic acid (CA) pathway included cholic acid, 3-oxo-cholic acid, 12-oxo-chenodeoxycholic acid, and deoxycholic acid. The main primary bile acids were defined as taurochenodexycholic acid, taurocholic acid, glycocholic acid, glycochenodeoxycholic acid, cholic acid and chenodeoxycholic acid and the main secondary bile acids as deoxycholic acid, glycodeoxycholic acid, taurodeoxycholic acid, glycolithocholic acid, lithocholic acid and taurolithocholic acid. Relative abundance values were calculated as a fraction of the total bile acids in each sample.

### DNA extraction and 16S rRNA analysis of the Afribiota stool samples

The microbiota profiles of the children included in Afribiota were described previously^31,32^. Briefly, gastrointestinal samples were extracted according to manufacturer’s instructions using commercial kits (QiaAmp cador Pathogen Mini or cador Pathogen 96 QIAcube HT Kit; Qiagen) with an additional bead-beating step to increase mechanical disruption. The efficiency of the extraction protocols was compared in between the two study sites using a Zymogen Mock Community. Extracted DNA was stored at -80 °C until sequencing. Extracted DNA samples were shipped to a commercial provider, where library generation and sequencing were performed (Microbiome Insights). Library preparation was performed as recommended by Kozich et al.^49^ using primers v4.SA501 to v4.SA508 and v4.SA701 to v4.SA712. The amplicon library was sequenced on a MiSeq using the MiSeq 500 Cycle V2 Reagent Kit (250 Å∼ 2). Data are available on the ENA server under accession number PRJEB48119. Demultiplexed reads were processed using the Dada2 pipeline^50^, and taxonomy was assigned using the Silva Database (version 128) as previously described^30^. The taxonomy table, metadata, and the ASV table for the samples used in this study can be found in Supplementary Tables S1-S3.

### In vitro screening of growth effects on isolated bacteria and synthetic communities

To test the effect of specific bile acids on members of the microbiome and especially bacteria associated with undernutrition and EED, we used a set of reference strains representing bacterial species that are either enriched or depleted in their relative abundance in the microbiome of children suffering of stunting and environmental enteric dysfunction (EED) (Table S1). The final set of bacteria tested included representatives of the oral microbiome, enteropathogenic species, which are both shown to be enriched groups within the gastrointestinal tract of stunted children^29,31,32,51^ and a set of (butyrate-producing) bacteria, which have been previously shown to be underrepresented in stunted children^28,32^. In total, 34 strains across 21 genera were tested for their growth phenotype in the presence of 19 different bile acids. Strains were tested under anaerobic (2% H_2_, 12% CO_2_, and 74 % N_2_) and microaerophilic conditions (5% O_2_, 10% CO_2_, and 85% N_2_) at 37°C and 42°C, respectively (specific conditions see Table S4) using a bile acid concentration of 0.25, 0.5 or 1 mM, consistent with standard high-throughput screening protocols^52^. Growth inhibition was measured over 22 hours, growth curves were truncated at the transition to exponential phase, and the AUC was calculated relative to the control (DMSO). Each strain was measured in at least three biological replicates. A significant inhibitory activity was defined as a false discovery rate (FDR) of <0.01. The raw AUC data can be found in Table S5 (single strains) and Table S6 (synthetic communities).

### Evaluation of bile acid effects on a synthetic gut microbial community in vitro

We selected 14 bacterial taxa commonly detected in the gut microbiota of stunted children and six underrepresented taxa (Supplementary Table S4). Strains were grown on BHI + blood agar plates under anaerobic conditions, and a single colony was then grown in 5 mL of BHI medium overnight. Cultures were combined in equimolar fractions to reach a total OD578 of 0.0125. A total of 500 µL of the prepared bacterial suspension was added to each well of a 96-deep-well plate pre-filled with 100 µL of BHI containing the bile acids to reach a final bacterial OD578 of 0.01 and bile acid concentrations of 1 mM, 500 µM, and 250 µM, respectively. To monitor growth kinetics, 100 µL of the community-bile acid mixture was transferred into a clear 96-well flat-bottom plate (Thermo Fisher Scientific) and monitored for growth as described above for the single strains. The deep-well plates with the remaining 500ul of bacteria-bile-acid mixtures were sealed with an AeraSeal breathable membrane (Sigma-Aldrich, cat. no. A9224) and incubated anaerobically at 37 °C for 24 hours. After incubation, 300 µL of culture was pelleted and cryopreserved for downstream 16S rRNA gene amplicon sequencing.

### 16S rRNA gene amplicon sequencing of the synthetic communities

Bacterial DNA was extracted from culture pellets using the DNeasy UltraClean 96 Microbial Kit (Qiagen, cat. no. 10196-4) or the DNeasy PowerSoil HTP 96 Kit (Qiagen, cat. no. 12955–4). Sequencing and library preparation were carried out by the NGS Competence Center NCCT (Tübingen, Germany). DNA concentration was measured using the Qubit dsDNA BR/HS Assay Kit (Thermo Fisher), and 100 ng of input DNA was used for library preparation. Final libraries were checked for fragment size using an E-Gel 96 Gel system with 2% mSYBR Safe DNA Gel Stain (Fisher Scientific), quantified via QuantiFluor dsDNA System (Promega), pooled equimolarly, and sequenced on an Illumina MiSeq with a v2 kit (2×250 bp reads, 10 pM input, 20% PhiX spike-in).

### Sequence processing and taxonomic assignment

Amplicon sequences were processed in R (v.4.2.0) using the dada2^50^ (v1.21.0) package, following the standard workflow. After inspecting quality profiles, reads were trimmed and filtered (trimLeft: 23/24; truncLen: 240/200; maxEE: 2; truncQ: 11), dereplicated, and ASVs inferred with default parameters. Paired reads were merged per sample, and merged reads were filtered by length (250–256 bp). Chimeras were removed. Taxonomic classification followed a two-step approach. First, ASVs were classified to the genus level using a curated DADA2-formatted version of the GTDB (release R06-RS202). Then, ASVs belonging to expected genera were further classified to species using a custom database containing full-length 16S rRNA sequences from the 19 community members. Species-level assignments were performed using the DECIPHER^53^ (v.2.24.0), with a threshold of >98% sequence identity to a known member. Abundance data were aggregated at the species level.

### Stool-derived in vitro communities and in vitro assays

To generate stool-derived *in vitro* communities (SICs), 0,2 g of frozen stool samples from African children (see Table S7) where resuspended in 3 mL of pre-reduced phosphate-buffered saline (PBS) in an anaerobic chamber filled with a gas mixture of 7% H_2_, 20% CO_2_ and 73% N_2_. Samples were let sit for 10 min at room temperature to allow large stool fragments to moist before vortexing them for 2 minutes and mixing them by pipetting up and down to disintegrate the particles before settling food particles for one minute again at room temperature. Subsequently, 50µL of the supernatant were transferred into 10 mL of pre-reduced BHI medium (Difco^Tm^ Brain Hearth Infusion, Becton Dickinson, Ref. 237500) supplemented with inulin from chicory at 1g/l (Sigma-Aldrich, cat. number I2255, Darmstadt, Germany), before incubation at 37°C. The inoculation was performed for all samples in three replicates. Every 48 hours, the SICs were passaged by transferring 5µL of the vortexed bacterial cultures in 1mL of fresh BHI-inulin medium in 96 deep well culture plates (Nunc® 96 DeepWell™ plate, non-treated, Merck, Cat. No. Z717274-60EA) at 1:200 dilutions and plates covered by a lid to avoid evaporation. A total of 5 passages were performed and after the last passage, 500µL of each stabilized SIC were thoroughly mixed with 500µL of pre-reduced glycerol (40%) containing palladium beads (Precipitated palladium, Sigma-Aldrich, Cat. No. 520810) and then stored at -80°C until needed. The sequencing data (ASV table, taxonomy table, metadata) of the initial fecal samples used for the SICs can be found in Supplementary Tables S8-S10. Frozen SICs were revived by inoculating the glycerol stocks in 1mL BHI-inulin in deep-well plates and allowing bacteria to grow for 48 hours in an anaerobic chamber, followed by one passage to re-stabilize the community. Just before conducting a treatment, the SICs grown for 48 hours were adjusted to an OD of 0.012 before the addition of the bile acid treatments. A bile acid treatment can be described as follows: 100µL of the 6x bile acid stock solutions was added to 500µL of the SIC cultures previously adjusted at O.D. 600 nm = 0.012. This resulted in a SIC culture at a starting O.D. 600nm = 0.01. The treated SICs were then incubated for 24 hours at 37°C under anaerobic conditions. DNA was extracted from a volume of 200µL as described below.

### DNA extraction and 16S sequencing of the SICs

At the end of the experiment, 100µL of SIC culture and 900µL of CTAB buffer (MaxWell RSC PureFood GMO and Autentification Kit, Promega, REF AS1600) were transferred into a QIAGEN Power Bead Pro Tube (QIAGEN, Cat. No. 19301). Tubes were incubated at 95 °C for 10 minutes and allowed to cool down for 2 minutes before performing lysis using a benchtop vortex and vortex adapter (QIAGEN, Cat. No. 13000-V1-24) for bead beating at speed 10, for 10 minutes. The samples were centrifuged at 14,000 x g for 10 minutes at room temperature. Then, 600µL of the supernatant was transferred into new 1,5 mL Eppendorf tubes. 40µL of Proteinase K and 20µL of RNase A (MaxWell RSC PureFood GMO and Autentification Kit, Promega, REF AS1600) were added to each sample, mixed by vortexing briefly and incubated at 70°C for 10 min. The final step of DNA extraction (DNA cleaning) was performed using the MaxWell RSC 48 apparatus (Promega, REF AS8500) using the “PureFood GMO protocol”. Purified DNA quality and quantity were assessed using the Nanodrop2000 system (Thermo Fisher Scientific) before sequencing. After DNA purification, 16S rRNA gene amplicon sequencing of the bacterial communities was performed by Novogene Co. Ltd. (United Kingdom) using 515F and 806R primers targeting the V4 regions of the 16S rRNA gene. Libraries were sequenced on an Illumina NovaSeq6000 apparatus (250pb paired end), with at least 30 000 raw sequences per sample. Quality control, including filtering, trimming, dereplication, chimera removal, and taxonomic assignment (Silva Reference Database version 128) of the reads was performed with the dada2^50^ (v.1.22.0) pipeline in R version 4.1.2. The raw data (taxonomy, ASV count and metadata table) can be found in Supplementary Tables S11-S13) The raw fastq files will be deposited on ENA under accession number (TBD).

### Culturing of reference strains for mass spectrometry analysis

*Shigella flexneri*, *Fusobacterium nucleatum*, *Streptococcus salivarius*, *Granulicatella elegans*, *Coprococcus comes* (now newly named *Bariatricus comes*), and *Roseburia intestinalis* type strains (Table 4) were selected as reference strains for mass spectrometry (MS) analysis. Strains were retrieved from frozen glycerol stocks and streaked onto pre-reduced BHI blood agar plates. After overnight incubation under anaerobic conditions (12% CO₂, 2% H₂, balance N₂) in a Coy anaerobic chamber, single colonies were inoculated into 5 mL of liquid medium for each strain as described in Table 4. Chenodeoxycholic acid, taurochenodeoxycholic acid, lithocholic acid, and taurolithocholic acid (Sigma-Aldrich) were dissolved in DMSO to prepare 50 mM stock solutions in V-bottom plates. Stock solutions were mixed with 50 µL of medium in U-bottom 96-well plates (Thermo Fisher Scientific) to yield final concentrations of 1 mM bile acid and 2% DMSO per well. Plates were transferred into the anaerobic chamber one day before inoculation to allow pre-reduction.

Overnight cultures were measured at OD₅₇₈ and diluted to a starting OD₅₇₈ of 0.02. A 50 µL aliquot of each diluted culture was added to the pre-reduced bile acid-containing wells and mixed, resulting in final concentrations of 0.5 mM bile acid and an initial OD₅₇₈ of 0.01. Plates were sealed with AeraSeal breathable membrane (Sigma-Aldrich, cat. no. A9224) and incubated at 37 °C for 24 h under anaerobic conditions.

Following incubation, plates were removed from the anaerobic chamber and centrifuged at 4,000 rpm for 5 min. Twenty microliters of supernatant from each well was transferred to a new U-bottom plate for subsequent MS analysis. All experiments were performed in at least three independent biological replicates.

### Mass spectrometry analysis

*Metabolite extraction.* Liquid samples (20 μL) were processed for LC–MS analysis using an organic solvent extraction method. In brief, a mixture of acetonitrile:methanol (1:1) was added to the samples (5:1 v/v). The mixtures were then incubated at -20°C for at least 1h and centrifuged at 4500 rpm for 15 min at 4°C. Supernatants were collected and subjected to two successive 30-fold dilutions with water before LC-MS analysis. *LC–MS analysis.* Analysis was performed by reversed-phase chromatography with n InfinityLab Poroshell HPH-C18 column (2.1×100 mm, 1.9 μm) and an Agilent 6546 LC/qTOF system. Water with 0.1% formic acid and acetonitrile with 0.1% formic acid was used as mobile phases A and B, respectively, with a gradient of 5 to 95% phase B in 5.4 min and a flow rate of 0.4 ml/min. The qTOF instrument was operated in negative scanning mode (100–1700 *m/z*) with source settings of 3,500 V for VCap, 2,000 V for nozzle voltage, a gas temperature of 225°C, a drying gas flow of 13 l/min, nebulizer pressure at 20 psig, and sheath gas settings of 225°C temperature and 12 l/min flow. Online mass calibration was performed using a second ionization source and a constant flow of reference solution (112.9856 and 1033.9881m/z, Agilent Ref. number G1969-85001). *Targeted feature detection (MS1).* The MassHunter Quantitative Analysis Software (Agilent, version 10.0) was used for peak detection and integration based and *m/z* and RT values of pure compound standards: chenodeoxycholic acid (m/z 391.2849, RT 4.17 min); taurochenodeoxycholic acid (m/z 498.2890, RT 3.37 min), lithocholic acid (m/z 375.2900, RT 4.88 min), taurolithocholic acid (m/z 482.2940, RT 4.13 min). *m/z* tolerance was set to 20 ppm, RT tolerance was set to 0.2 min.

### Microscopy analysis of Roseburia intestinalis and Coprococcus comes exposed to different bile acids

To assess the effects of bile acids on butyrate-producing gut bacteria, strains were cultured under anaerobic conditions and analyzed for growth kinetics and cell morphology. Strains were streaked on BHI + blood agar plates and incubated anaerobically. Strains were inoculated in 5 mL PYI medium and grown overnight at 37 °C, followed by a second overnight subculture under the same conditions. The second culture was diluted to a final OD578 of 0.01 in a total volume of 100 µL containing 500 µM bile acids in U-bottom 96-well plates. For microscopic phenotyping, 0.3 µL of mid-exponential phase cells was spotted onto a 1% (w/v) agarose pad. Images were acquired using a Nikon Eclipse Ti2-E microscope equipped with a monochrome Nikon DS-Qi2 camera and a CFI P-Apo DM 100x Lambda Oil Ph3 objective. Image segmentation was performed in ImageJ v2.16.0 using the Trainable Weka Segmentation plugin (https://doi.org/10.1093/bioinformatics/btx180, v4.0.0). Segmentation-derived morphology features and CFU data were analysed in R v4.5.1.

### Statistical analyses

Statistical analysis of bile acid data was performed in R version 4.1.2 as reported in the accompanying R Markdown file deposited under https://github.com/VonaeschLabUNIL/Afribiota/tree/main/Bile_acids. Batch effect was corrected using ComBat from the sva package^54^ (v.3.45.0), which uses an empirical Bayes framework to model and adjust for known batch effects. ComBat correction was applied to log-transformed bile acid abundance data individually within each sample type (fecal, gastric and duodenal), with country and stunting given as biological outcomes of interest (to account for expected differences in the data) and mass spectrometry batch as the effect to be corrected for. Bile acids that were less than 20% prevalent within their compartment were also filtered out. ComBat-corrected data was used for PCA ordination of country and stunting, and for nonparametric tests (Wilcoxon and Spearman’s correlation) of differences by country, HAZ and stunting, and AAT. Multivariate linear models were performed on log-adjusted, prevalence-filtered, but combat-uncorrected bile acids data, controlling for batch as a covariate. Models for country adjusted for batch, sex, age, haz, inflammation (measured through calprotectin and AAT levels in the feces), and anemia. Models for HAZ or stunting adjusted for batch, country, age, sex and inflammation (measured through calprotectin and AAT levels in the feces). Models for cfu count in the duodenum (as a measure of SIBO) where performed, controlling for age, batch and sex and with an interaction term for anemia and country of origin. Multivariate models for inflammation markers were performed on the continuous variables and controlling for batch, country, age, sex, stunting status. Finally, multivariate models for associations with hemoglobin were performed controlling for batch, age, sex, inflammation and with an interaction term for country of origin and haz-score within the duodenal space was controlled for batch, age, sex, country of origin and inflammation within the small intestine (calprotectin and AAT levels in the duodenum). Statistical analyses and visualizations of the microbial data, bacterial isolates, synthetic communities and stool-derived *in vitro* communities were conducted in R version 4.4.2 using phyloseq^55,56^ (v.1.46.0), MicroEco^57^ (v.0.9.19), vegan^58^ (v.2.6.8), microbiomeMarker^57^ (v.1.8.8), ape^59^ (v.5.8.1) and ggplot2^60^ (v.3.5.1) packages. The envfit function from the vegan package was used to assess which environmental variables explained the clustering of the samples by fitting environmental vectors (bacterial taxa relative abundance, bile acid levels and other factors such as age, sex, country or sequencing run) on the ordination space of a Principal Coordinates Analysis (PCoA) based on Bray-Curtis distance. Spearman correlation was used to assess the relationship between bile acid levels and bacterial taxa relative abundance, and statistical significance was adjusted using Benjamini–Hochberg correction. Multivariate analyses of differentially abundant taxa were performed on pooled samples from both countries as well as on data from each country independently. Chi2 test using Bonferroni correction for multiple testing was used to assess for the presence/absence of bacterial taxa in the samples in the upper and lower quantiles for the lithocholic acids and derivatives. Differential abundance analysis of the bacterial taxa in the upper and lower quantiles group, as well as in the stool-derived *in vitro* communities, was performed with LefSe from the microbiomeMarker^61^ package using a LDA cutoff of 2 and a Benjamini–Hochberg correction. Alpha diversity was assessed using the estimate_richness function from the phyloseq package using the observed and Shannon indexes, and groups were compared using the Wilcoxon rank-sum test. Absolute abundance was calculated by multiplying the relative abundance by the DNA concentration measured in the samples after DNA extraction. Log2 fold change with a pseudo-count of half the minimum value was used to assess for the change in taxa relative abundance within the synthetic communities treated with bile acids.

For the analysis of MS1 targeted data, extracted peak areas were analyzed to determine whether a compound was depleted in samples treated with the corresponding compound. Outliers were identified and then removed using Tukey’s method, which defines outliers as values below Q1 − 1.5×IQR or above Q3 + 1.5×IQR, where Q1 and Q3 are the 25th and 75th percentiles, and IQR is the interquartile range (Q3 − Q1). Pairwise comparisons of peak areas between experimental groups (control, bacterial culture, and supernatant) were performed using two-sample t-tests. Resulting p-values were adjusted for multiple comparisons using the Benjamini-Hochberg false discovery rate (FDR) correction. Peak areas were normalized by scaling to the maximum mean value within each compound-media group. Significant compound degradation or modification was defined using an FDR threshold of ≤ 0.05. Second, hydrolysis products of taurine-conjugated acids were manually identified and assessed in sample groups where metabolite levels increased. Pairwise t-tests were used to compare metabolite levels between control, bacterial culture, and supernatant samples, and p-values were then adjusted for multiple comparisons using the Benjamini-Hochberg FDR correction. Peak areas were normalized by scaling to the maximum mean value within each compound-media group. Significant compound degradation or modification was defined using a p-value of ≤ 0.05.

## Results

### Bile acid profiles differ along the gastrointestinal tract and between the two study sites

We performed bile-acid-targeted metabolomics on gastric (N=216), duodenal (N=226), and fecal (N=935) samples from children in Madagascar and the Central African Republic (CAR) (Supplementary Figure S1). Fecal samples were available from both stunted- and non-stunted children, whereas gastric and duodenal samples were collected via aspiration from stunted children only.

As expected, there were drastic differences in bile acid composition throughout the intestinal tract (Figure 1A, PERMANOVA p<0.001). Upper intestinal samples (gastric and duodenal) were generally similar to one another but had distinct bile acid composition compared to fecal samples. Upper intestinal samples were dominated by the primary bile acids glycocholic acid and glycochenodeoxycholic acid, while fecal samples were dominated by the secondary bile acids deoxycholic acid and lithocholic acid (Figure 1B). Primary and conjugated bile acids were therefore significantly enriched in upper intestinal samples, whereas secondary and unconjugated bile acids were enriched in feces, consistent with bile acid metabolism by colonic gut bacteria (Figure 1C-E, Kruskal-Wallis p<2e-16 for all using either raw or batch-corrected values). Together, these data confirm the reliability of both the sample type and bile acid measurements collected in this study.

**Fig 1.**
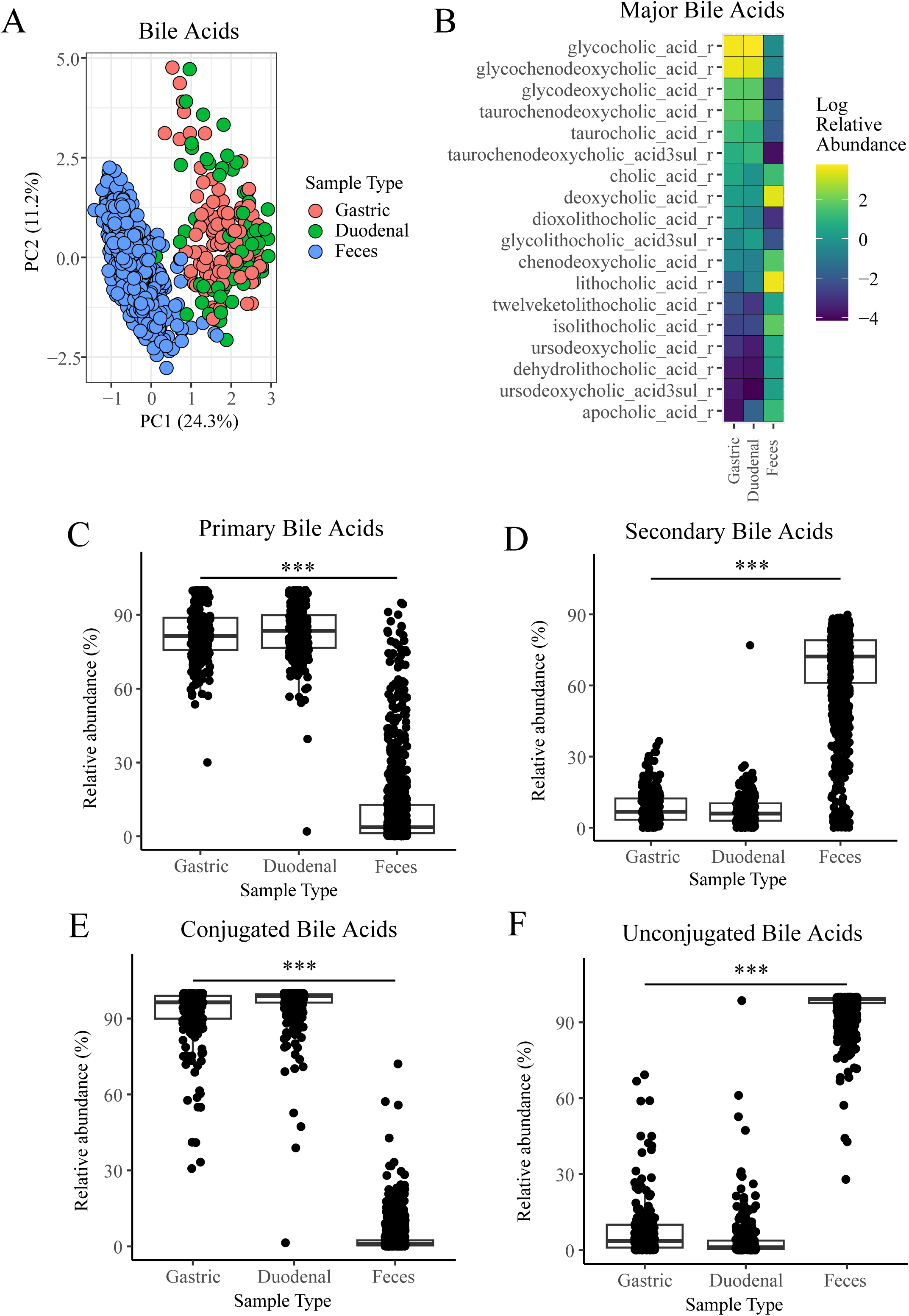
Differences in bile acids along the child’s gastrointestinal tract. (A) PCA ordination of bile acids by sample type. (B) Topmost abundant bile acids (≥1% mean relative abundance) by sample type. (C-F) Relative abundance of the major bile acid categories by sample type: (C) primary, (D) secondary, (E) conjugated, and (F) unconjugated bile acids. Significance determined by PERMANOVA (A) or by Kruskal-Wallis test (C-F).

Metabolomic profiling was performed in three distinct batches due to the large number of samples; this led to a visible batch effect in addition to the dramatic effect of sample type (Figure 1A, Supplementary Figure S2). We corrected the batch effect within sample type using ComBat (see methods and Supplementary Figure S2) for subsequent analyses.

A minor difference in bile acid profile by country was detected in fecal samples, but not in upper intestinal samples (PERMANOVA, duodenum; p=0.156, gastric, p=0.155; feces, p=0.001, Supplementary Figure S3A-C). Fecal samples from children in CAR had fewer primary bile acids remaining in feces, and a greater abundance of conjugated bile acids (Wilcoxon p=0.008 and p=7.446e-06, respectively, Supplementary Figure S3D-E). Primary bile acids remained significantly different by country in linear models adjusting for age, sex, stunting, inflammation, and anemia; however, conjugated bile acids were not different by country after adjustment for co-variates, suggesting that one of these co-variates might be contributing to country-specific differences. Furthermore, several specific bile acids had a differential relative abundance in the fecal samples according to the country of origin of the children, and many of these remained significant in adjusted linear models (Supplementary Figure S3F).

We have previously described differences in the intestinal microbiome composition between these two study sites^31,32^, which might influence the production of secondary bile acids; conversely, an altered bile acid milieu might influence microbiome composition. In summary, bile acid composition differs drastically along the gastrointestinal tract and modestly in feces between countries.

### Lithocholic acids are decreased in the feces of children with growth stunting

We have previously shown that stunted children in both Madagascar and CAR have a distinct microbiome composition characterized by an enrichment of oral bacterial species in feces^31,32^. We therefore investigated whether bile acid composition also differs by stunting status. To test this, we ran a combination of FDR-corrected nonparametric tests and FDR-corrected linear models that accounted for batch effect, country, age, sex, and inflammation (fecal calprotectin and alpha-1-antitrypsin (AAT)).

Interestingly, using an FDR-corrected Wilcoxon test, we found that lithocholic acid and related bile acids (lithocholic acid derivatives; see definition in Methods) were significantly decreased in the feces of stunted children (Figure 2A-B). This association was also significant in an FDR-corrected Spearman’s association against continuous height-for-age z-score (HAZ) (Figure 2C), and by raw p-value in a linear model with covariates (Supplementary Table S14). A handful of other bile acids, including ursocholic acid and derivatives (7 beta-hydroxy epimer, combining ursocholic acids and ursodeoxycholic acids and their derivatives) as well as cholic acid and derivatives, were increased in stunted children by linear model (Figure 2D & E, Supplementary Table S14). There was no difference in the abundance of these bile acids by stunting severity in the duodenal or gastric samples, although no non-stunted controls are available for the duodenum. Nevertheless, these data suggest a difference in bile acid processing by colonic bacteria during undernutrition, which might further influence the gut microbiome or the host.

**Fig 2.**
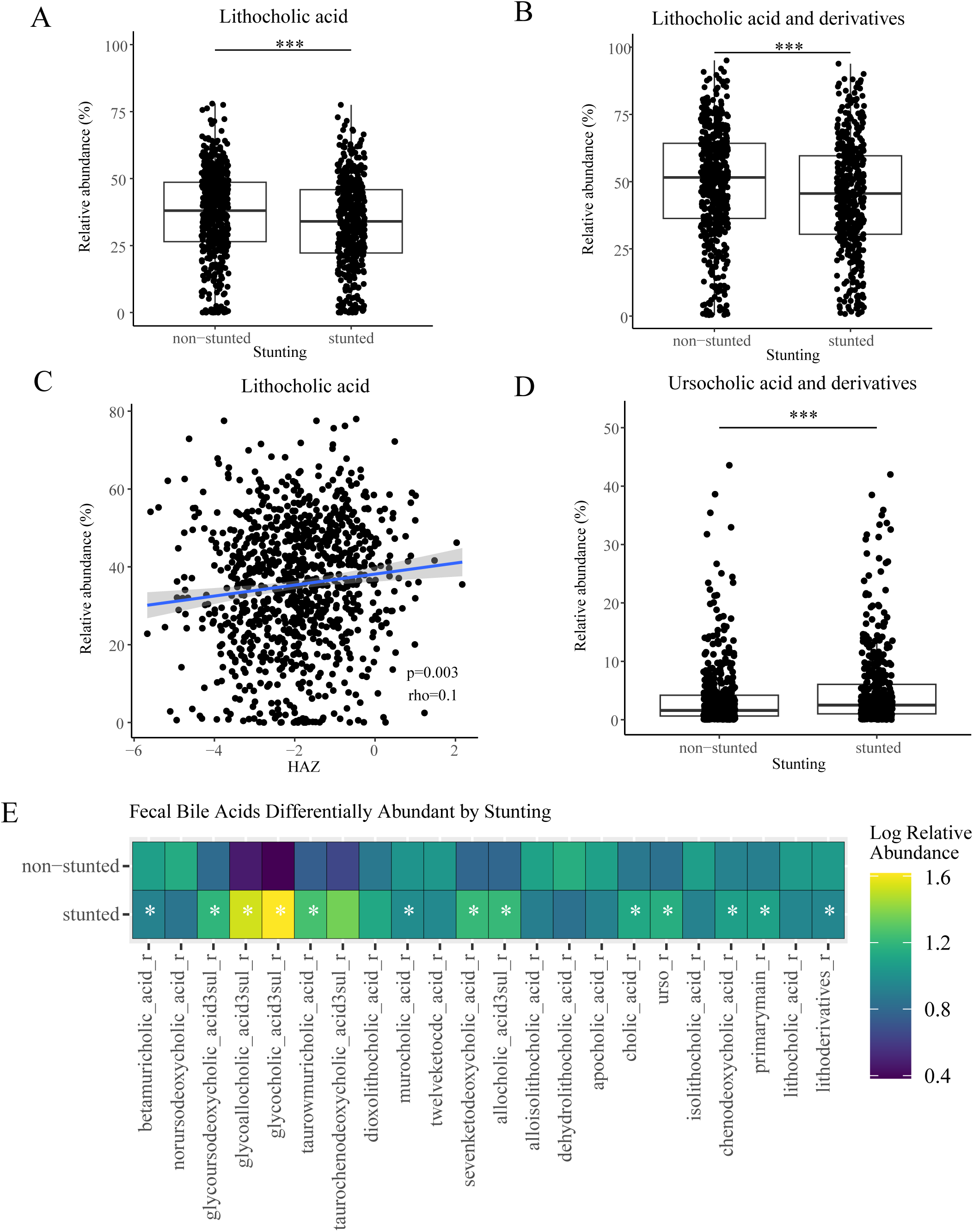
Differences in fecal bile acids by stunting status. (A) Fecal lithocholic acid and (B) all fecal lithocholic acid derivatives, by child stunting status. (C) Lithocholic acid relative abundance in feces by height-for-age z-score (HAZ). (D) Fecal ursocholic acid and derivatives by child stunting status. (E) Abundance of all individual bile acids by stunting in feces, which were significant according to FDR-corrected Wilcoxon tests on batch-adjusted data. A (*) indicates that significance was maintained in a linear model correcting for batch, country, age, sex, and gastrointestinal inflammation.

### Biomarkers of EED are associated with profound changes in the overall bile acid pool of the feces

Stunting is often accompanied by environmental enteric disorder (EED), an inflammatory condition of the intestinal tract, and we have shown before that there are independent changes in the microbiome induced by stunting and EED^31^. To test for fecal bile acids associated with the EED markers alpha-1-antitrypsin (AAT) or with the presence of small intestinal oral bacterial overgrowth (SIOBO), we ran a combination of Benjamini-Hochberg (BH)-corrected nonparametric tests and BH-corrected linear models that accounted for batch effect, age, sex, country of origin, HAZ and calprotectin levels (Table S14). We observed few convincing associations with SIOBO (Table S14). Consistent with the stunting patterns observed, however, higher inflammation as measured by AAT was modestly associated with decreased fecal levels of lithocholic acid derivatives and secondary bile acids, and increased levels of primary bile acids (Figure 3A-C); these trends were significant by FDR-corrected non-parametric test, but not in corrected linear models. Interestingly, AAT levels were positively associated with total sulfated bile acids, and to a lesser extent with ursocholic acid derivatives, according to both FDR-corrected nonparametric tests and FDR-corrected linear models (Figure 3D-E & Table S14). These data imply a strong association of bile acid pools with EED markers, suggesting an additive effect of undernutrition and EED on the overall pool of bile acids.

**Fig 3.**
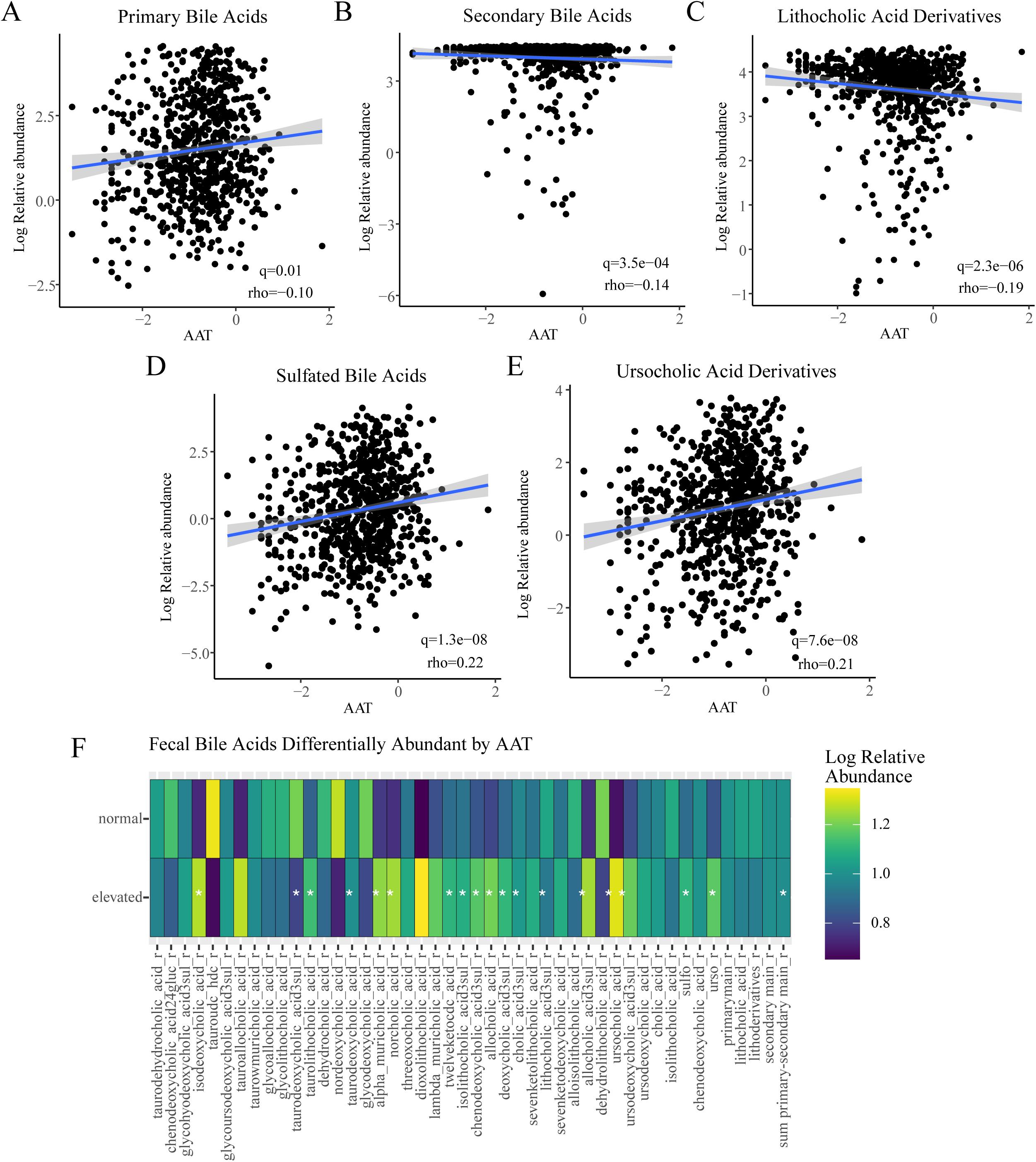
Differences in fecal bile acids by fecal alpha-1-antitrypsin (AAT). The fecal abundances of (A) primary bile acids, (B) secondary bile acids, (C) lithocholic acid and derivatives, (D) sulfated bile acids, and (E) ursocholic acid and derivatives, compared to fecal AAT levels. Significance was determined by FDR-corrected Spearman’s correlations on batch-adjusted data. Regression lines are drawn by a simple linear fit. (F) Abundance of all individual bile acids by AAT category in feces, which were significant according to FDR-corrected Spearman’s correlations against the continuous variable on batch-adjusted data. A (*) indicates that significance was maintained in a linear model correcting for batch, country, age, sex, calprotectin, and height-for-age z-score. ATT levels are given in log scale.

### Secondary bile acids in the feces are correlated with the abundance of EED-related bacteria

It is well known that members of the commensal microbiota can convert primary into secondary bile acids and chemically diversify the main bile acids into numerous bile acid derivatives observed in the intestinal tract. Leveraging microbiome data available from previous studies in Afribiota, we thus aimed to assess whether bile acid levels in the fecal samples associated with the presence of any bacterial taxa.

We observe that overall clustering of the samples on a bacterial PCoA plot is mostly associated with the relative abundance of *Prevotella* 9 (envfit: R^2^ = 0.82, p-value < 0.05), followed by *Bacteroides* (envfit: R^2^ = 0.28, p-value < 0.05) (Table S15). Conversely, bile acids show weaker associations with the observed diversity. Using envfit, we could show that the bile acids associated most strongly with beta diversity include alloisolithocholic acid (R^2^ = 0.11, p-value < 0.05), isolithocholic acid (R^2^ = 0.11, p-value < 0.05) and lithocholic acids and derivatives (R^2^ = 0.10, p-value < 0.05). Taken together, secondary bile acids (R^2^ = 0.08, p-value < 0.05) and primary bile acids (R^2^ = 0.04, p-value < 0.05) show modest associations with the ordination. Of note, bile acids were more strongly associated with the overall composition than other factors such as country (R^2^ = 0.02, p-value < 0.05), age of the children in years (R^2^ = 0.02, p-value < 0.05), or sequencing run (R^2^ = 0.06, p-value < 0.05).

We next assessed the relationship between specific genera and bile acids using Spearman correlation, corrected with Benjamini-Hochberg (BH) (Table S16). Notably, several genera of the *Ruminococcaceae family*, *Anaerotruncus* and a member of the *Eubacterium coprostanoligenes* group, which have all members that are known butyrate producers, were significantly associated with secondary bile acid levels (Figure 4A). This association of butyrate-producers with secondary bile acids and notably lithocholic acid and derivatives was also observed independently in both study settings (Supplementary Figure S4A-B, Table S17). Furthermore, by selecting the upper and lower quantiles for the values of lithocholic acids and derivatives (170 samples each in the upper and lower quartiles, respectively) and by assessing for the presence/absence of genera within each group using a Chi2 test (Bonferroni-corrected p-values < 0.05), we found 39 genera to be differentially present (Table S17). Using LefSe to analyze these relationships, we identified several genera differentially abundant in the two groups (LDA > 2 and BH-corrected p-values < 0.05) (Figure 4B), of which the circles indicate the taxa that were also significant on the Chi2 test (38/39 taxa). Of note, several bacteria of oropharyngeal origin, including *Streptococcus*, *Haemophilus* and *Fusobacterium* as well as the possible enteropathogens *Escherichia/Shigella,* among others are enriched in samples low in lithocholic acid and derivatives and several known butyrate-producing bacteria, including several members of the *Ruminococcaceae* and *Lachnospiraceae* families, *Anaerotruncus*, member of the *Eubacterium coprostanoligenes* group, *Butyricimonas* and *Desulfovibrio* among others, are enriched in the group with high levels of lithocholic acid and derivatives (Figure 4B). Several of these taxa show a linear relationship with the bile acids (Figure 4C-H, Supplementary Figure S4C-M).

**Fig 4.**
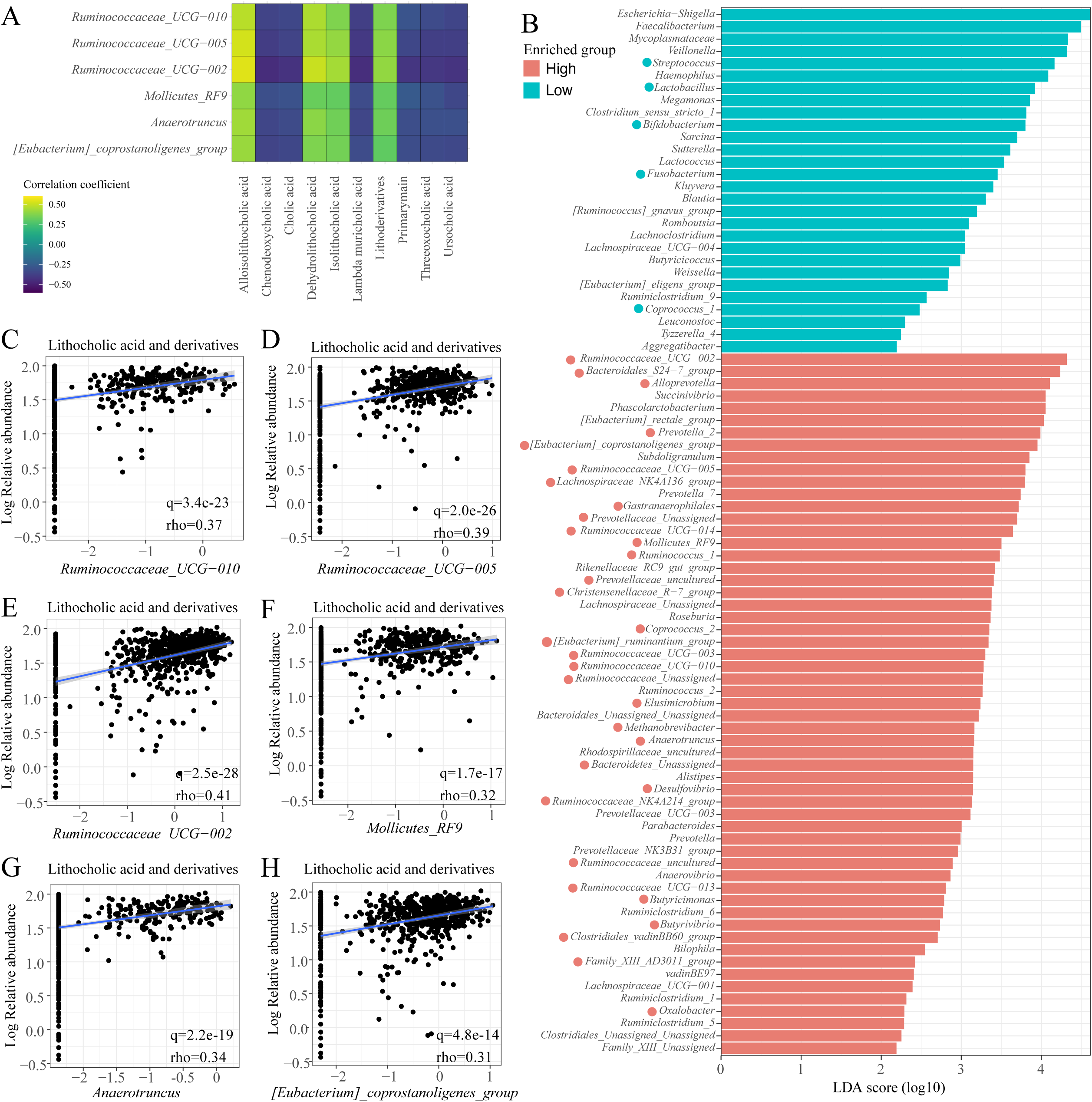
Correlation of bile acid levels with bacterial taxa in the feces. (A) Heatmap of the strongest correlations between bile acids and bacterial genera in the Afribiota samples (all q.values < 0.05). (B) Lefse LDA score plot of the bacterial genera enriched in the high and low lithocholic acid samples. Dots show the genera differentially present/absent by Chi2 test in the high (uppest quartile) and low (lowest quartile) lithocholic acid samples. (C-H) Linear relationship between the six bacterial genera with the strongest correlation and the relative abundance of lithocholic acid and derivatives in fecal samples.

In conclusion, there are clear associations between given bile acids and given bacterial taxa, especially a positive association between butyrate-producing bacteria and a negative association between oro-pharyngeal taxa and secondary bile acids, suggesting that there might be a direct interaction in between these two entities, at least in specific cases.

### Secondary bile acids boost the growth of butyrate-producing bacteria in vitro and in fecal explants

Although colonic bacteria process bile acids, they are also directly affected by the bile acid milieu. We hypothesized that altered bile acid composition in the feces of stunted children might exacerbate microbiome differences. We therefore tested whether secondary bile acids affected the growth of a panel of human gut bacteria using a semi-high-throughput screening set-up described previously^60^ (Figure 5A, Supplementary Table S18). Interestingly, we found that bile acids affect the oro-pharyngeal taxa significantly more than butyrate-producing bacteria (Figure 5B & D). Furthermore, lithocholic acid and derivatives seem not only to spare butyrate-producing bacteria but even promote their growth based on their growth curves as measured by OD (AUC > 1) (Figure 5C & E).

**Fig 5.**
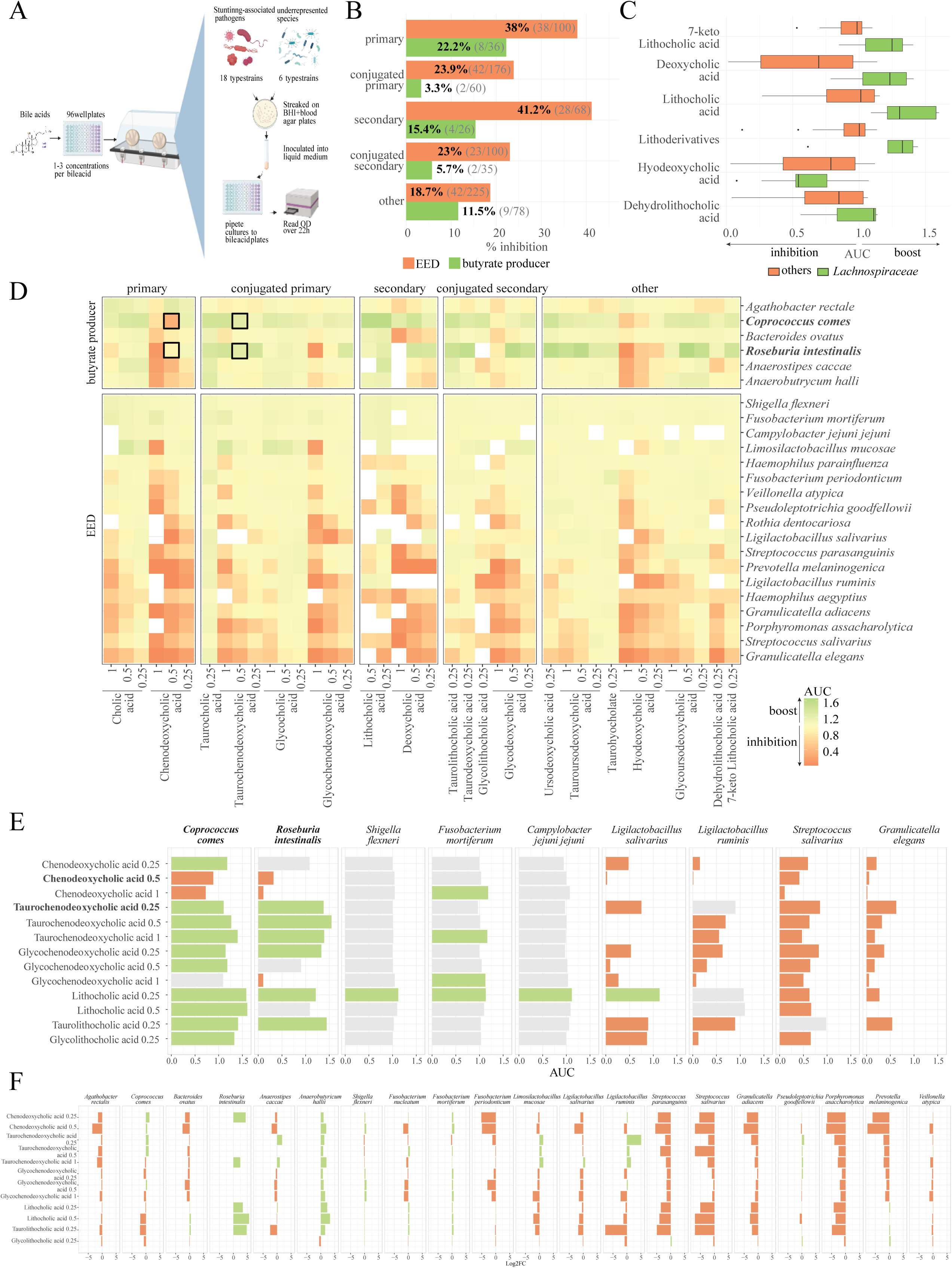
Secondary bile acids boost butyrate producers *in vitro*. (A) Overview of *in vitro* bile acid testing on EED-associated bacteria. A panel of 19 bile acids was tested for their effects on the growth of 18 type strains of stunting-associated pathogens and six underrepresented commensal strains. (B) Effect of specific bile acids on the growth of health-associated and EED-associated taxa. Bar plots and bold numbers indicate the percentage of inhibitory effects of bile acids on bacterial growth in each category (orange: EED-associated taxa, green: butyrate producers/health-associated taxa). Absolute counts are shown in brackets (number of negative growth outcomes / total bile acids tested). (C) Box plots displaying the area under the curve (AUC) for *Lachnospiraceae* versus other taxa across different bile acids. An AUC above 1 indicates increased growth relative to the control (DMSO). Whiskers represent the most extreme data points within 1.5× the interquartile range (IQR), with outliers shown individually. (D) Heatmap showing the normalized area under the growth curve (AUC) for a panel of bile acids tested at different concentrations on health-associated and EED-associated type strains. AUC values are normalized to the respective solvent-treated controls. Each tile represents the mean of 2 to 4 biological replicates. (E) AUC for a panel of bile acids tested at different concentrations on butyrate producers and oro-pharyngeal type strains is shown in (D). An AUC value above 1 (green) indicates increased growth relative to the control (DMSO treatment). An AUC value below 1 (orange) indicates decreased growth relative to the control (DMSO treatment). (F) Log2 fold change (Log2FC) of the relative abundance of the type strains in the synthetic community treated with a panel of bile acids. Log2FC above 0 (green) indicates increased growth relative to the control (DMSO). Log2FC below 0 (orange) indicates a decreased growth relative to the control (DMSO).

To validate these findings in independent experiments, we focused on two butyrate-producing species, *C. comes* (now called *Bariatricus comes*) and *R. intestinalis*, and a selected panel of bile acids. The inhibitory effect of chenodeoxycholic acid at 0.5 mM (highlighted in Figure 5D & E) was confirmed for both species (Supplementary Figure S5A). For taurochenodeoxycholic acid, which was predicted from the screen to increase the area under the growth curve (AUC, highlighted in Supplementary Figure S5D-E), we were able to reproduce this effect. In *C. comes*, the increase was observed when applying a cutoff after the exponential phase, whereas in *R. intestinalis*, it occurred at the transition into the exponential phase (Supplementary Figure S5B). Taurolithocholic acid, which was expected to increase AUC for both species at 0.25 mM based on the screening data, showed a concentration-dependent effect: the increase was abolished at higher concentrations (0.5 mM) in both species (Supplementary Figure 5C). This highlights the strong concentration dependence of bile acid effects. Consistent with this, bile acids also modulated cell size in a species- and compound-specific manner at the cellular level (Supplementary Figure S5D & E).

To determine whether bile acid sensitivity is linked to direct metabolism or intracellular accumulation, we systematically tested these possibilities for selected bile acids and species (Supplementary Tables S19-S26, Figures 6A & C). Bile acids that increased AUC in specific species (*i.e.,* taurochenodeoxycholic acid, taurolithocholic acid) appeared to be bioaccumulated and deconjugated by those species (*R. intestinalis, C. comes*) (Figures 6C & D), suggesting a potential detoxification mechanism. In contrast, bioaccumulation patterns of lithocholic acid in *G. elegans* (Figure 6A&B) as well as chenodeoxycholic acid in *F. nucleatum* (Supplementary Figure 6) might explain the toxic effect of these compounds in these oral taxa.

**Fig 6.**
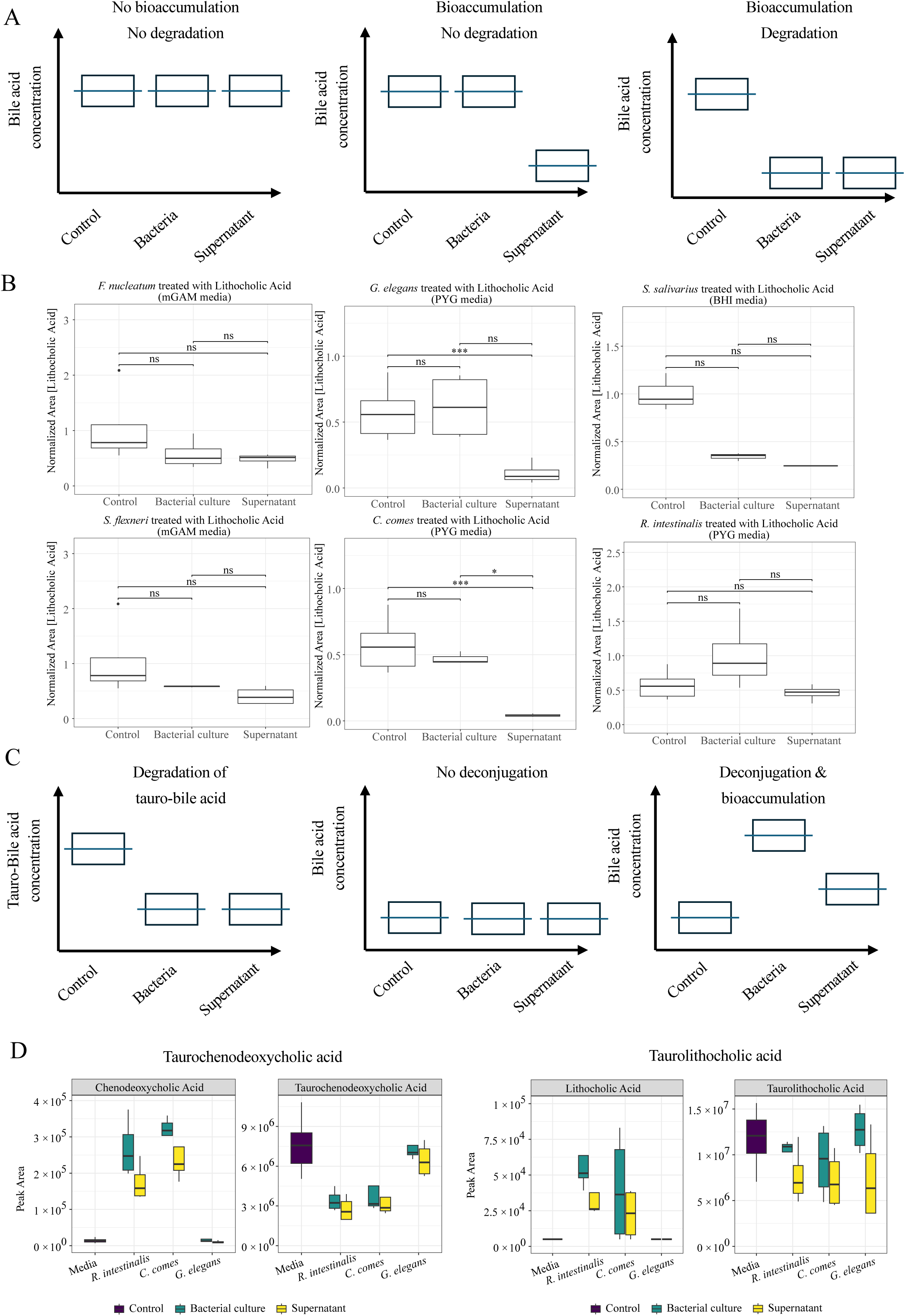
Bile acids are bioaccumulated and deconjugated by given oral and butyrate-producing taxa. (A) Expected levels of bile acids in the experimental set-up in the context of bioaccumulation and/or bile acid degradation. (B) Normalized lithocholic acid concentration in the medium control, bacterial culture or supernatant after overnight growth of *F. nucleatum, G. elegans, S. salivarius, S. flexneri, C. comes* or *R. intestinalis* in the presence of this bile acid. (C) Expected levels of bile acids in the context of degradation and deconjugated of tauro-conjugated bile acids. (D) Measurement of Taurochenodeoxycholic acid and Chenodeoxycholic acid, Taurolithocholic acid and lithocholic acid in the medium incubated with the tauro-conjugated forms of these bile acids as well as *R. intestinalis* or *C. comes*. *: FDR<0.05; ** FDR<0.01; *** FDR<0.001 in a two-sample t-tests. Resulting p-values were adjusted for multiple comparisons using the Benjamini-Hochberg false discovery rate (FDR) correction.

Last, enteropathogens, as expected from previous literature (reviewed in^62^), do not seem to be affected in their growth by bile acids (AUC ∼ 1) (Figure 5D & E). These associations were also seen in a synthetic community composed of the different reference strains (Figure 5F, Table S4). Most oro-pharyngeal taxa showed decreased abundance in the presence of bile acids, while butyrate producers such as *Roseburia intestinalis* and *Anaerobutyricum hallii* increased in abundance in the presence of most bile acids tested (Figure 5F).

To confirm these associations in complex multi-species communities, we established stool-derived *in vitro* communities (SICs) derived from fecal samples of children from the Afribiota project (Figure 7A). In total, ten SICs were established in three replicates, of which five samples were from the Central African Republic (two normally nourished, three undernourished) and five from Madagascar (three normally nourished, two undernourished). While there were expected overall shifts in the community composition between feces and SICs, the SICs represented many strains of the original samples, with some taxa increasing in relative abundance, while others decreased/disappeared (Figures 7B & C and Supplementary Figure S7). Several families were conserved in the SICs, including the *Bacteroidaceae*, *Butyricicoccaceae*, *Clostridiaceae*, *Coriobacteriaceae*, *Eggerthellaceae*, *Enterobacteriaceae*, *Enterococcaceae*, *Lachnospiraceae*, *Oscillospiraceae*, *Peptostreptococcaceae*, *Ruminococcaceae* and *Veillonellaceae*.

**Fig 7.**
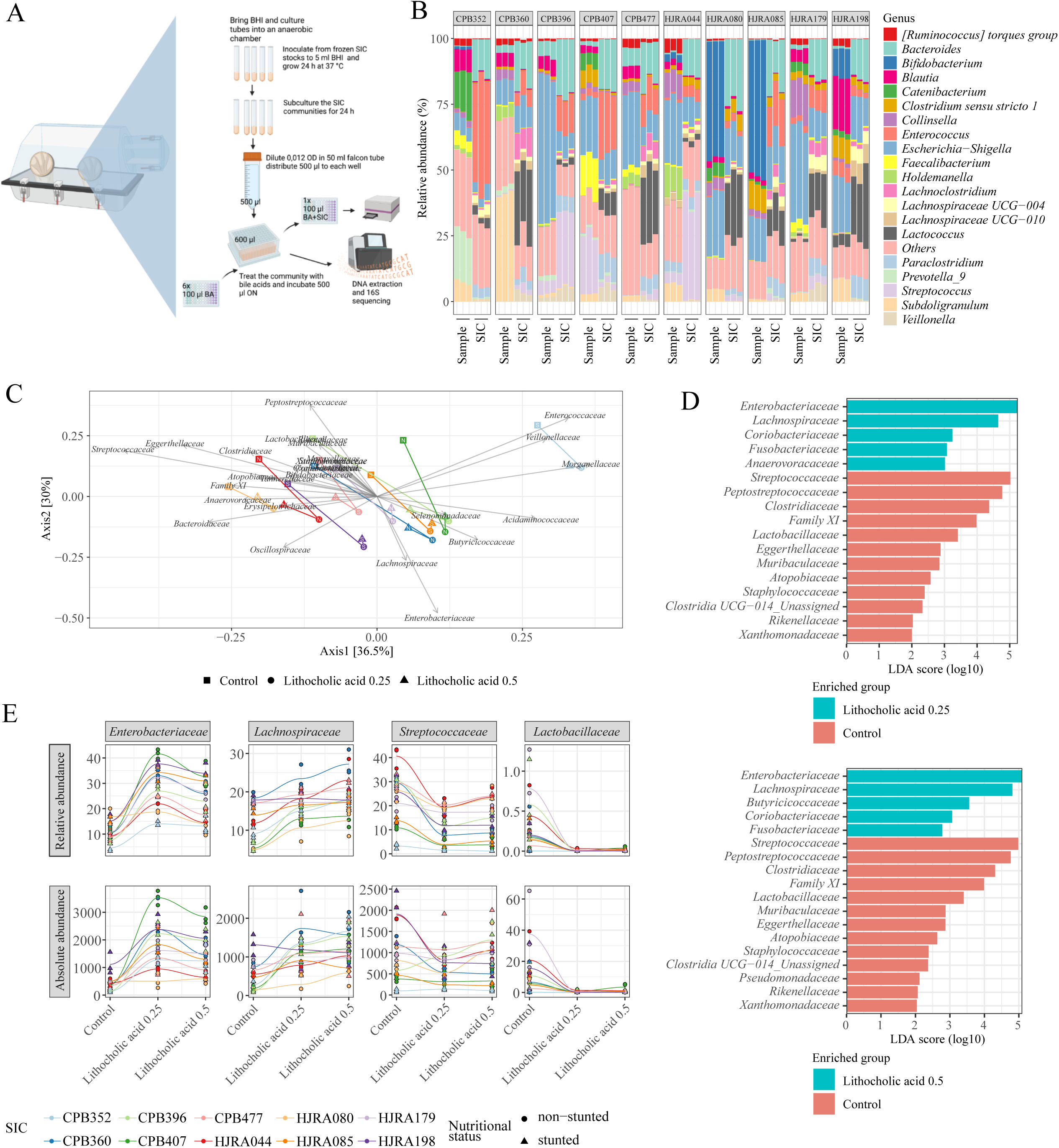
Lithocholic acid boosts butyrate producers in full fecal communities. (A) Overview of *in vitro* bile acids testing on Stool-derived *in vitro* Communities (SICs). Ten stabilized SICs (5 from Central African Republic including 2 normally nourished, 3 undernourished, 5 from Madagascar including 3 normally nourished, 2 undernourished) were resuspended in brain-heart infusion medium (BHI) and then transferred into deep multi-well plates containing the different bile acids. (B) Relative abundance of the top 20 most abundant genera comparing the fecal samples with their respective stabilized SICs. The Others category regroup all the less abundant genera. (C) Principal Coordinate Analysis (PCoA) based on Bray-Curtis distance at the family level of the SICs treated with lithocholic acid. Each point represents the centroid of the three SIC replicates, shapes represent the different treatment conditions. SIC from stunted children are depicted with a S inside the shapes. SIC from non-stunted children are depicted with a N inside the shapes. Arrows represent the direction and strength of the significant correlation between bacterial families and the ordination space as identified by envfit (p<0.05). (D) Bacterial families enriched in the control or SIC treated with lithocholic acid at two different concentrations as identified by LefSe (LDA > 2 and q-value<0.05). (E) Visualization of the changes in the relative (upper panel) and absolute (lower panel) abundance of *Enterobacteriaceae, Lachnospiraceae, Streptococcaceae* and *Lactobacillaceae* upon treatment with lithocholic acid at two different concentrations. Absolute abundances were estimated by multiplying the relative abundance by the DNA concentration of the extracted SICs.

Once stabilized the SICs were treated with different concentrations of the secondary bile acids lithocholic acid, taurolithocholic acid, and the primary bile acids chenodeoxycholic acid and taurochenodeoxycholic acid. Overall alpha diversity changes upon bile acid treatment were small, yet largely consistent across the different SICs, with some exceptions, especially regarding the Shannon index (Supplementary Figure S8). Furthermore, the addition of the bile acids led to clear shifts in the overall microbiome composition that were largely conserved across the different SICs (Figure 7C). Tauro-conjugated bile acids showed a less consistent community shift compared to unconjugated bile acids (Supplementary Figures S9-S12). Most interestingly, we observed changes in several bacterial taxa upon treatment with the bile acids (Figure 7D & E, Supplementary Figure S9-S12). Lithocholic acid notably led to an increase in the relative abundance of *Lachnospiraceae* (envfit: R^2^ = 0.26, p-value < 0.05) and *Enterobacteriaceae* (envfit: R^2^ = 0.88, p-value < 0.05) and a relative decrease in *Streptococcaceae* (envfit: R^2^ = 0.82, p-value < 0.05), among others, mirroring previous findings from the Afribiota cohort and the single-strain exposures (Figure 7D). The same trends were observed in all SICs by both relative or absolute abundances (Figure 7E). Similar trends were observed for the treatment with chenodeoxycholic acid, but *Streptococcaceae* were unaffected by conjugated bile acids (Supplementary Figure S9-S12). Therefore, (secondary) bile acids have a positive effect on the growth of healthy human gut commensals while restricting several oropharyngeal taxa.

## Discussion

In this study, we show that lithocholic acid and its derivatives are decreased in the feces of children suffering from stunted growth. Furthermore, we show a decrease in lithocholic acids and derivatives in children with signs of EED. The levels of lithocholic acids in the feces are positively correlated with butyrate-producing bacteria and negatively associated with bacteria of oro-pharyngeal origin. Furthermore, we show a boosting effect of lithocholic acid and other secondary bile acids on butyrate-producing strains and the inhibitory effect on oro-pharyngeal bacteria *in vitro* in fecal explants, mirroring findings from the Afribiota cohort. The inhibitory effect on oral bacteria is also visible on isolated strains, while the growth-promoting effect on butyrate-producers seems to be a community effect, most likely induced through a stronger effect of the bile acids on the oral bacteria and the fact that these two bacterial groups show niche overlap^63^. Therefore, the suppression of the oral bacteria might lead to an indirect growth advantage of the butyrate-producers in the complex communities. Furthermore, we show that several butyrate-producers can accumulate and deconjugate bile acids, yet do not seem to degrade them. Jointly, our data suggest that secondary bile acids have a positive and direct effect on the growth of butyrate-producing gut commensals while restricting several oro-pharyngeal taxa.

Previous work on bile acid profiles in undernourished children showed a negative correlation of serum glycocholic acid with linear growth and a positive correlation of glycocholic acid in the duodenum and serum with biomarkers of inflammation and EED severity in children aged 3-6 and 9 months^40^. Furthermore, the same study showed lower levels of secondary bile acids in stunted children in both serum and duodenal samples. The authors also found a positive association between the levels of ursodeoxycholic acid, deoxycholic acid and glycoursodeoxycholic acid and a negative correlation with the levels of glycocholic acid and growth at 24 months^40^. In another study measuring bile acids in serum, the ratio of conjugated to unconjugated bile acids were negatively associated with HAZ^41^. Similar results were also obtained in the fecal samples collected from a mouse model of undernutrition^64^. Our data thus confirms the decrease of secondary bile acids and the increase in cholic acid and derivatives in stunted children compared to non-stunted controls, even though our data was measured in the feces, compared to the serum in the study by Zhang et al. and Moreau et al. However, we failed to see any association between stunting and duodenal bile acids, possibly also because we did not include non-stunted controls.

Previous work in mice further showed a decrease of taurocholic acid in the feces of undernourished mice suffering from EED^65^. While we did not find a significant association between taurocholic acid and stunting, there was a significant association between taurocholic acid and the fecal levels of alpha-1-antitrypsin (AAT), a marker of intestinal inflammation often used to define EED, even though the association was lost when correcting for multiple testing. Taurocholic acid was also associated with microbiota maturation in the intestinal tract of neonatal mice^66^, and the decrease in taurocholic acid in undernutrition could therefore be one of the factors contributing to the observed microbiota immaturity in young children suffering from undernutrition^67^.

Secondary bile acids are furthermore also decreased in a syndrome closely related to EED, inflammatory bowel disease (IBD), as shown in several studies in human subjects^68,69^. In these studies, there was likewise a positive correlation between *Ruminococcaceae* and secondary bile acids, mirroring clinical findings from our study and showing the relevance of our findings beyond the undernutrition field.

Secondary bile acids are closely associated with health. Previous studies show that lithocholic acid has beneficial effects on aging that are similar to those of calorie restriction. Lithocholic acid activates AMP-activated protein kinase (AMPK) with a beneficial effect on lifespan and health in Drosophila and *Caenorhabditis elegans*^70^. Therefore, some of the pathophysiological changes observed in undernutrition could, at least in part, be a direct effect of this secondary bile acid deficiency. Furthermore, there is clear indication that bile acids can regulate the virulence of several enteropathogens (reviewed in^62^), as demonstrated for example by *Vibrio cholerae*, where the hydrolysis of taurocholate to cholate by members of the intestinal microbiome blocks cholera colonization^71^. Changes in the pool of bile acids could thus also explain changes in virulence that might be associated with microbiome alterations in the context of childhood undernutrition and, for example, the high burden of asymptomatic colonization by enteropathogens^30^.

In our study, stunted/EED children further showed a significant increase in 7-α-hydroxymers (ursocholic acid and derivatives) compared to non-stunted/non-inflamed controls. 7-α-hydroxymers are generated through bile conversion by specific commensals, i.e. by *Clostridium absonum*^72^, *Clostridium barati*^73^ or *Ruminococcus gnavus* N53^74^. Nevertheless, little is known about these bacteria producing ursodeoxycholic acid and derivatives as well as the role of ursodeoxycholic acid and other 7-α-in childhood undernutrition. There is thus clearly more research needed in this field.

Stunted child growth was furthermore associated in our study with an increase in sulfated bile acids. Bile acid sulfation is mediated by a group of enzymes called sulfotransferases (SULTs) and is generally believed to be a pathway to detoxify and eliminate bile acids from the body through urine. Previous research has shown that they are also elevated in cholestatic diseases, and it was suggested that sulfation might contribute to the regulation of the overall bile acid pool under pathological conditions (reviewed in^75^). Most interestingly, there is also an increase of sulfated bile acids in the context of IBD, a closely related syndrome to EED (reviewed in^12^). The human microbiota can affect the pool of sulfated bile acids through a process called desulfation, with genera including *Clostridium*, *Peptococcus* and *Fusobacterium* harboring taxa that are able to perform this enzymatic reaction (reviewed in^76^). Of note, our study is the first to assess sulfated bile acids in undernourished children and children suffering from EED.

As mentioned before, in our clinical data, we saw a positive correlation between secondary bile acids and several members of the *Ruminococcaceae* and *Lachnospiraceae*. In fecal explants, administration of secondary bile acids, including lithocholic acid, notably led to an increase in the relative abundance of *Lachnospiraceae* and *Enterobacteriaceae* and a relative decrease in *Streptococcaceae*. Experiments with isolated strains furthermore showed that several butyrate producers can grow better in the presence of several secondary bile acids, suggesting that the bile acids and/or derived metabolites might be used as an additional energy source by these taxa. In line with this suggestion, it was shown previously that several commensals can bioaccumulate bile acids^77^. To date, most studies have tested for the bile-converting ability of bacteria yet have not considered secondary bile acids as an actual energy source. A recent article assessed the effect of different bile acids on a set of intestinal bacteria and found growth-promoting effects of ursodeoxycholic acid and tauroursodeoxycholic acid on the growth of *Bifidobacterium pseudonumeratum* and *B. pseudopodium*^77^. To the best of our knowledge, this is the only other report suggesting that the microbiota can thrive on secondary bile acids.

In mice, it has been previously shown that several gram-positive bacteria, normally found in the oral cavity or the duodenum, grow to higher abundances if the production of bile is blocked in the mice or humans. The most affected taxa are *Streptococcus thermophilus, Lactobacillus (para)casei* and *Lactococcus lactis*^78^, probably due to an absence of bile acid de-conjugating enzymes in these taxa. In the same study, these taxa also showed inhibition by glycochenodeoxycholic acid and glycocholic acid *in vitro*. Susceptibility to deoxycholic acid was also found in the pathogen *Streptococcus pneumoniae*^78^. Molecular studies showed that autolysins do not play a role in the toxic effect of bile acids but did not reveal the actual killing mechanism^77^. In lactobacilli and bifidobacteria it was suggested that cholic acid, deoxycholic acid, and chenodeoxycholic acid can damage membranes as well as DNA, cause oxidative and pH stress, and may chelate calcium and/or other cellular ions, altogether killing the bacteria^79^. In a recent *in vitro* screen, it was shown that deoxycholic acid can inhibit the growth of several gram-positive bacteria (*Streptococcus thermophilus* and *Lactobacillus rhamnosus*), which are similar to the oral bacteria we tested^77^. While the authors show that the effect of bile acid on bacterial growth can be dependent on overall media composition in some cases, the effect on these two gram-positive bacteria was medium-independent.

Overall, a more extensive characterization of the growth-promoting and killing effect of different bile acids on commensals of the gastrointestinal tract would allow for a better understanding of the bile acid-microbiome-host physiology triad.

Finally, our data showed a neutral effect of bile acids on single isolates of *Enterobacteriaceae*, yet a clear increase in their relative and absolute abundance in the context of fecal explants treated with secondary bile acids. This data is, at least to some extent, mirroring the results obtained in a previous study, where the authors observed a relative increase in *Proteobacteria* upon treatment with several secondary bile acids^77^. This observation could be linked to the formation of biofilms, as it was described previously that bile acids can induce biofilm formation in *Enterobacteriaceae*^80^ or simply to the competitive ability of *Enterobacteriaceae* to fill empty niches within a disturbed microbial ecosystem. Therefore, there seems to be higher-order interactions governing the overall community structure upon bile acid treatment and clearly more work is needed to understand the interactions between bile acids and the human intestinal microbiome in detail.

Our study has some limitations: due to ethical constraints, we were not able to collect duodenal samples from non-stunted children, thus lowering the power to detect differences in the pool of bile acids within the small intestinal content. Furthermore, as the microbiome analysis was performed using amplicon sequencing, our dataset provides no insight into the genetic capacity of the microbiome to enzymatically process bile acids. Despite these limitations, our study also has several strengths: first and foremost, our study includes a clinical study with almost 1000 children from two distinct geographical sites, and additionally uses experimental models to infer causality of the observed associations. To the best of our knowledge, our study is also the first to show the growth-promoting effect of secondary bile acids on butyrate-producing taxa, thus opening the way for further studies aimed at boosting these important players within the gastrointestinal tract and overall host physiology.

In conclusion, secondary bile acids lead to a clear decrease in bacteria of oral origin, while boosting the growth of several butyrate-producing taxa. Given the microbiome-modulating potential of several secondary bile acids, these observations might lay the groundwork for further studies aimed to use these findings for better treatments against childhood undernutrition targeting the microbiome. Such future therapies should be combined with other compounds that suppress the observed increase in Proteobacteria to ensure efficient community modulation.

## Author Contributions

Conceptualization: P.V., L. M. and P.J.S; methodology: P.V., K.H., N.A., J.R.G., L.M.; validation: M.W., K.H.; formal analysis: P.V., K.H., M.W., J.R.G., S.Y., N.A, J.H., C. G., L. B.; investigation: P.V., J.R.G., K.H., J.H., N.A., C. G., L. B.; data curation: P.V., K.H., M.W., J.R.G., P.V., K.H.; writing-original draft preparation: P.V., K.H.; writing-review and editing: all authors; visualization, P.V., K.H., J.R.G., L.B., N.A., S.Y.; supervision: P.V., L.M., B.B.F., C.H.B., P.J.S.: project administration: P.V.; funding acquisition: P.V., L.M., P.J.S. All authors have read and agreed to the published version of the manuscript.

## Supplementary Figures

**Fig S1.**
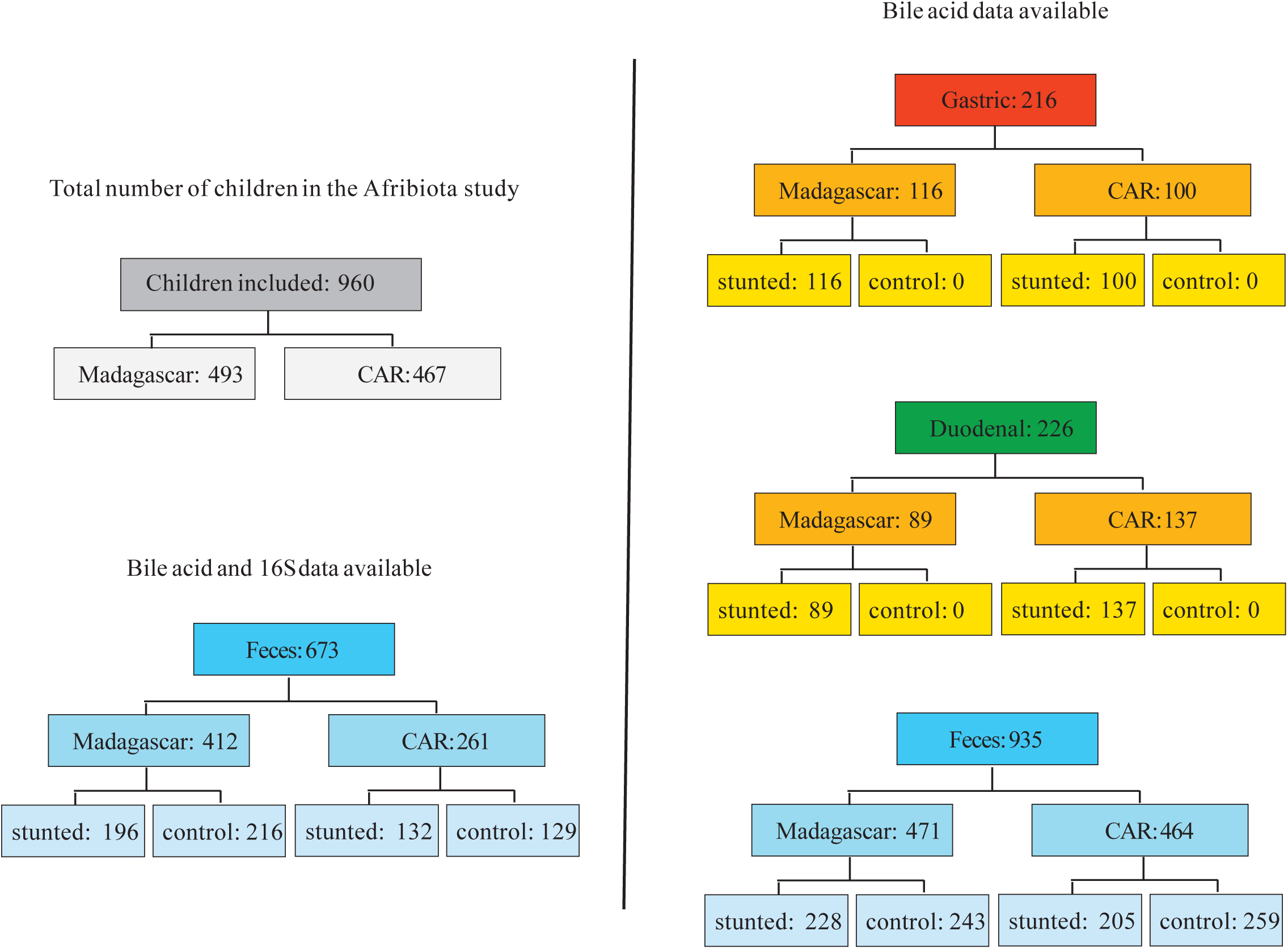
Flowchart showing sample size per compartment for all microbiome and bile acids data used in the current manuscript.

**Fig S2.**
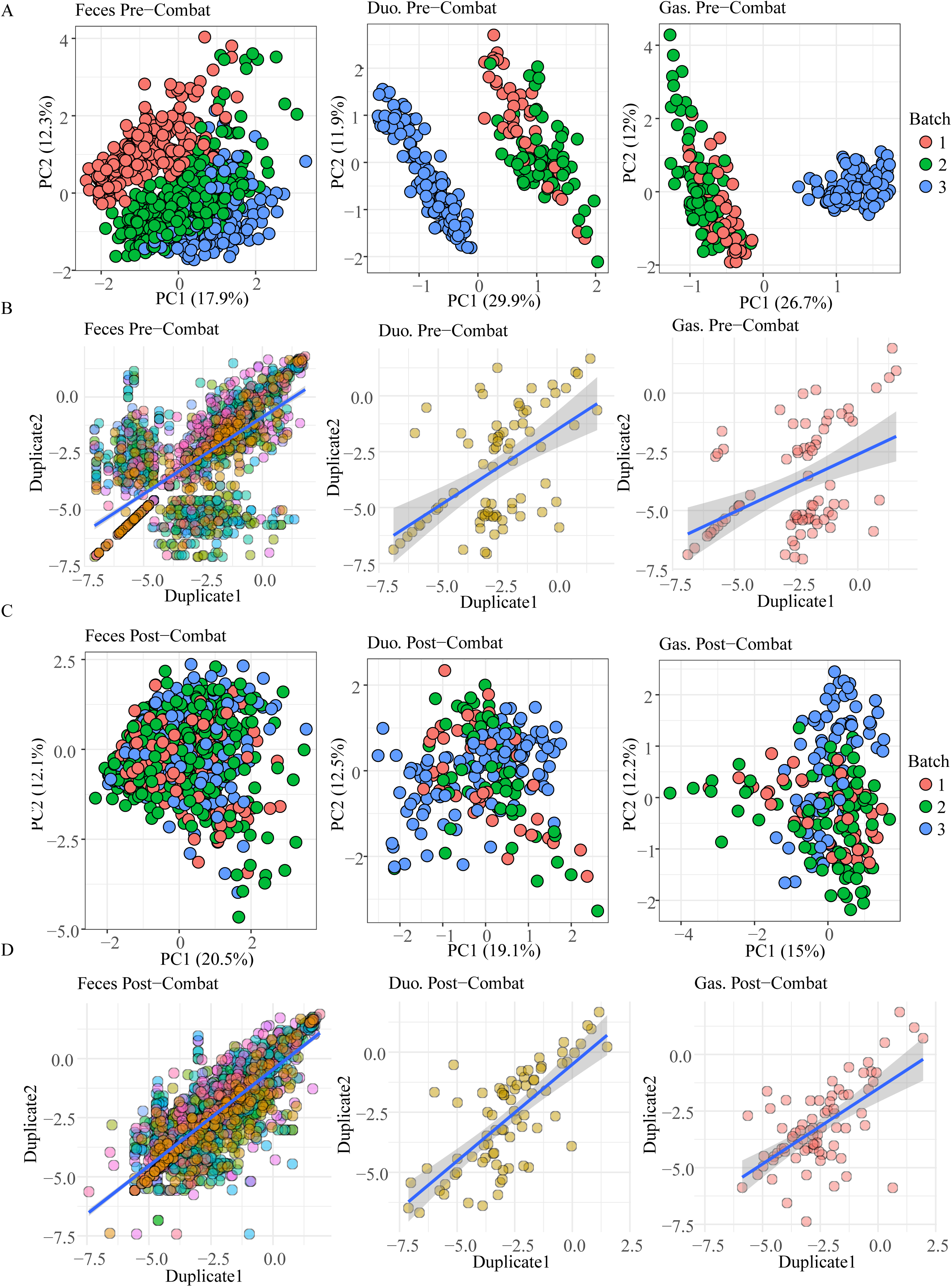
Batch effects and corrections. (A) PCA ordination of fecal, gastric and duodenal bile acids by mass spectrometry batch. (B) Correlation between technical duplicates (samples from the same participant processed twice for metabolomics) in fecal, gastric and duodenal compartments, prior to any batch correction, for all detected bile acids. Each data point is one person. (C) PCA ordination of fecal, gastric and duodenal bile acids by mass spectrometry batch, after ComBat-based batch correction. (D) Correlation between technical duplicates for all detected bile acids, following batch correction. Regression lines are determined by simple linear fit (B, D).

**Fig S3.**
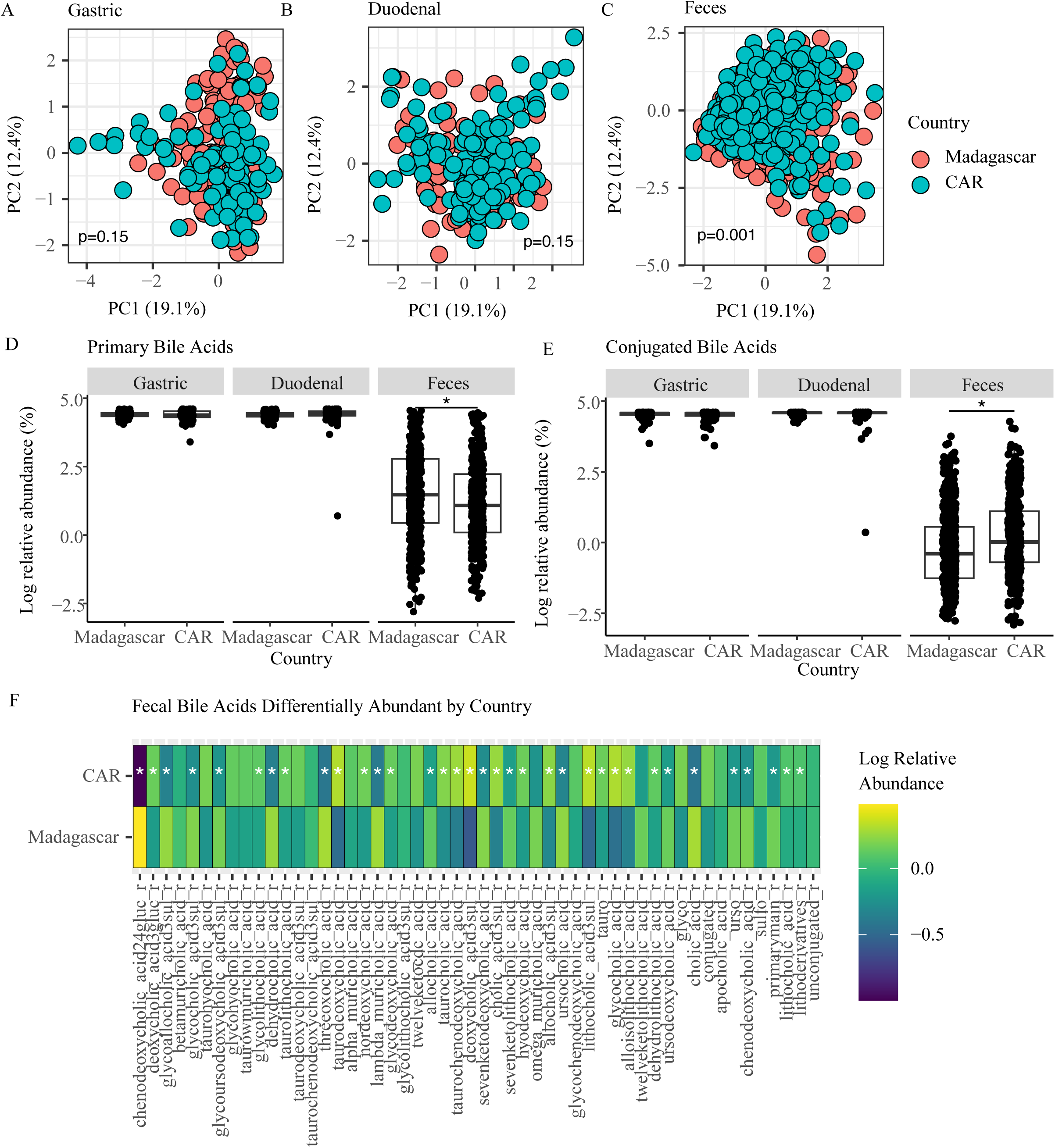
Differences in bile acids by country of origin. (A-C) PCA ordination of bile acids by country of origin in the (A) gastric, (B) duodenal and (C) fecal compartments. (D) Relative abundance of primary bile acids by country and sample type. (E) Relative abundance of conjugated bile acids by country and sample type. Significance determined by FDR-corrected Wilcoxon on batch-adjusted data (D-E). (F) Abundance of all individual bile acids by country in feces which were significant according to FDR-corrected Wilcoxon tests on batch-adjusted data. A (*) indicates that significance was maintained in a linear model correcting for batch, age, sex, haz, gastrointestinal inflammation and anemia.

**Fig S4.**
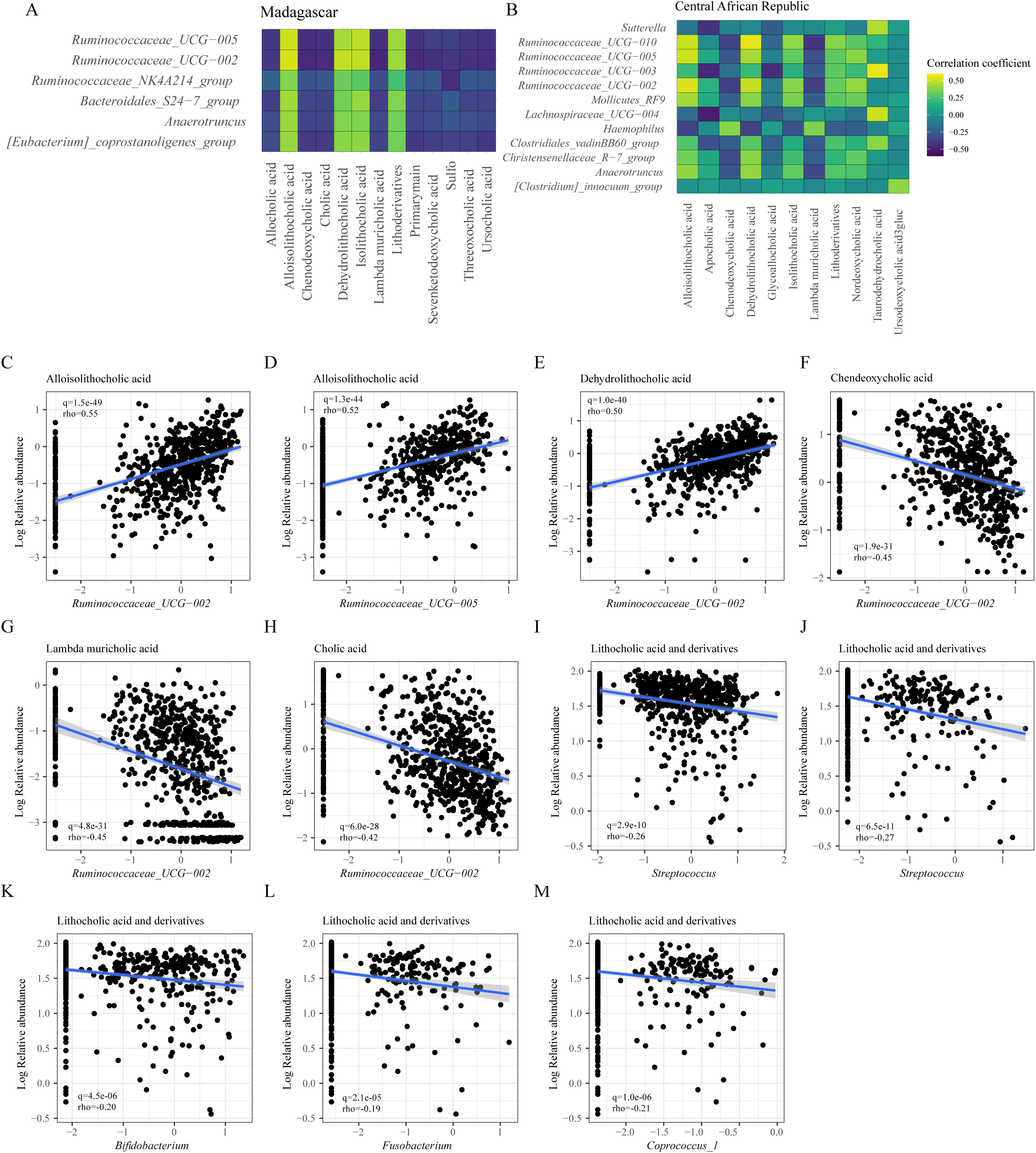
Bacteria-Bile acid correlations plots. (A) Heatmap of correlations between bacterial genera and bile acid levels in samples from Madagascar (all q.values < 0.05). (B) Heatmap of correlations between bacterial genera and bile acids levels in samples from the Central African Republic (all q.values < 0.05). (C-E) Linear relationship of the 3 strongest positive correlations found in the whole dataset. (F-H) Linear relationship of the 3 strongest negative correlations found in the whole dataset. (I-J) Linear relationship of the oral associated bacteria with lithocholic acid and derivatives.

**Fig S5.**
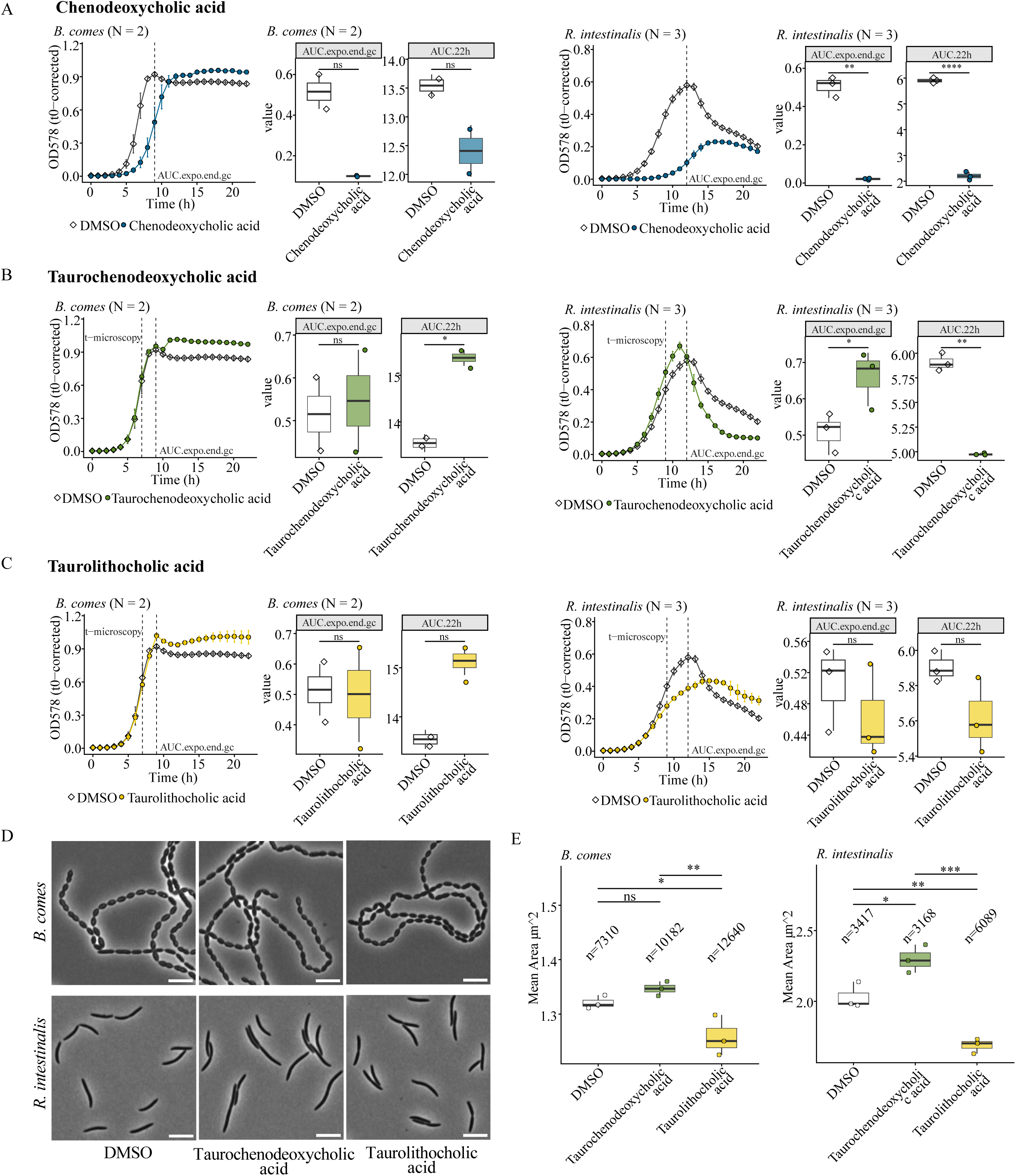
Bile acids modulate *R. intestinalis* and *C. comes* growth. (A) – (C) Growth curve analysis of *C. comes* and *R. intestinalis* treated with bile acids at 500 µM. “t-microscopy” indicates mid-exponential time point for microscopic phenotyping. “AUC.expo.end.gc” indicates the time point for AUC calculation at the end of the exponential phase of the DMSO growth control. Pairwise tests with one-way ANOVA followed by Benjamini-Hochberg adjustment for multiple comparisons adjusted p-values < 0.05 (*), < 0.01 (**), < 0.001 (***), < 0.0001 (****), ns: not significant. (D) Microscopic phenotyping of mid-exponential phase *C. comes* (7h) and *R. intestinalis* (9h) treated with 500 µM bile acids. Representative phase contrast images of treated bacterial species. scale = 5 µm. N = 3. (E) Cell size segmented phase contrast images. One-way ANOVA followed by Tukey’s HSD post hoc test for multiple comparisons adjusted p-values < 0.05 (*), < 0.01 (**), < 0.001 (***), ns: not significant. n = number of cells from 3 technical replicates with 3 fields of view each. N = 3.

**Figure S6.**
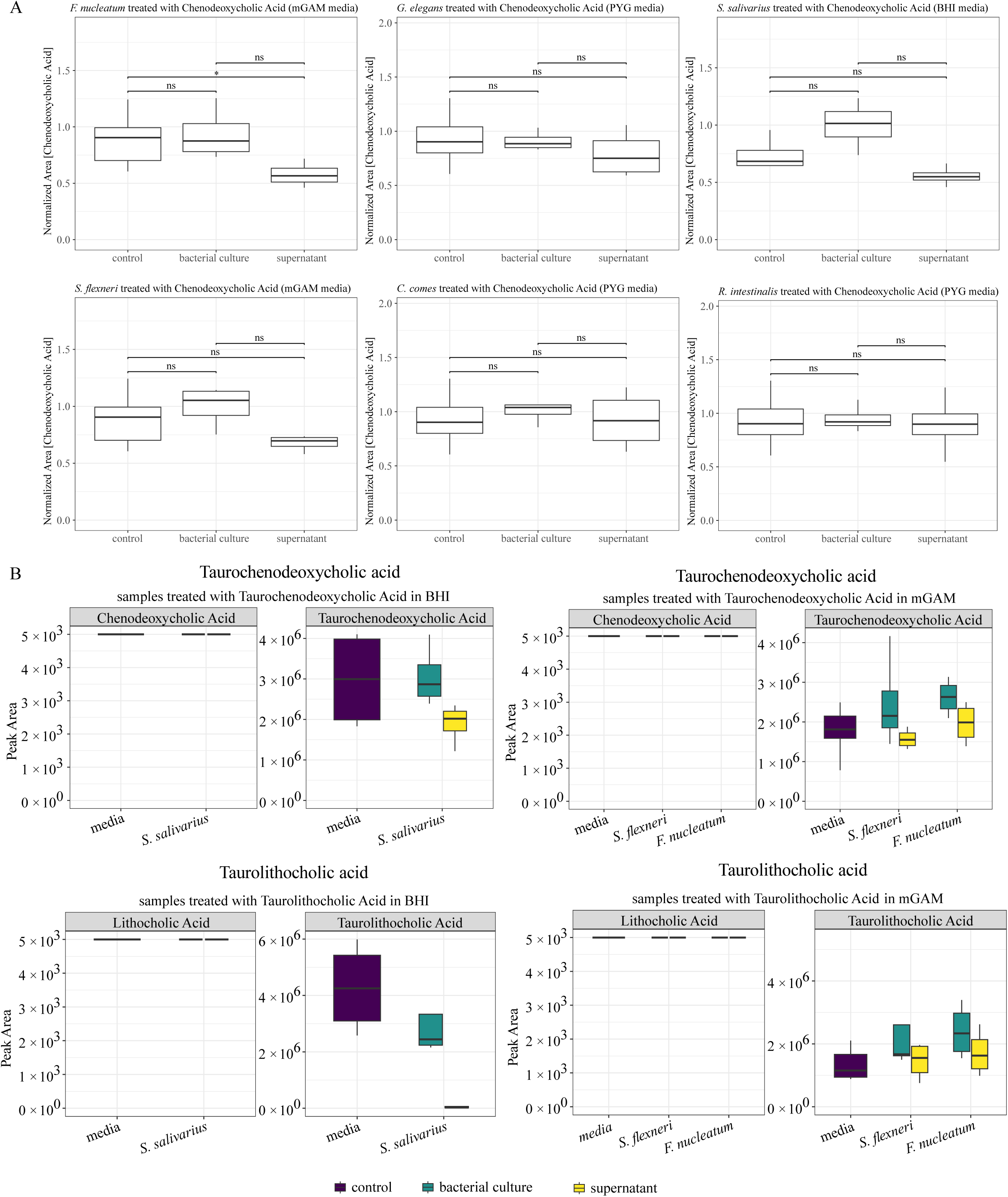
Bile acid bioaccumulation and deconjugation in a set of reference bacteria. (A) Normalized area of Chenodeoxycholic acid in the medium, as well as in the cell pellet and supernatant of cells grown overnight in medium supplemented with the respective bile acids (B) Measurement of Taurochenodeoxycholic acid and Chenodeoxycholic acid and Taurolithocholic acid and lithocholic acid in the medium incubated with the tauro-conjugated forms of these bile acids as well as *S. salivarius, S. flexneri* or *F. nucleatum*. *: FDR<0.05; ** FDR<0.01; *** FDR<0.001 in a two-sample t-tests. Resulting p-values were adjusted for multiple comparisons using the Benjamini-Hochberg false discovery rate (FDR) correction.

**Fig S7:**
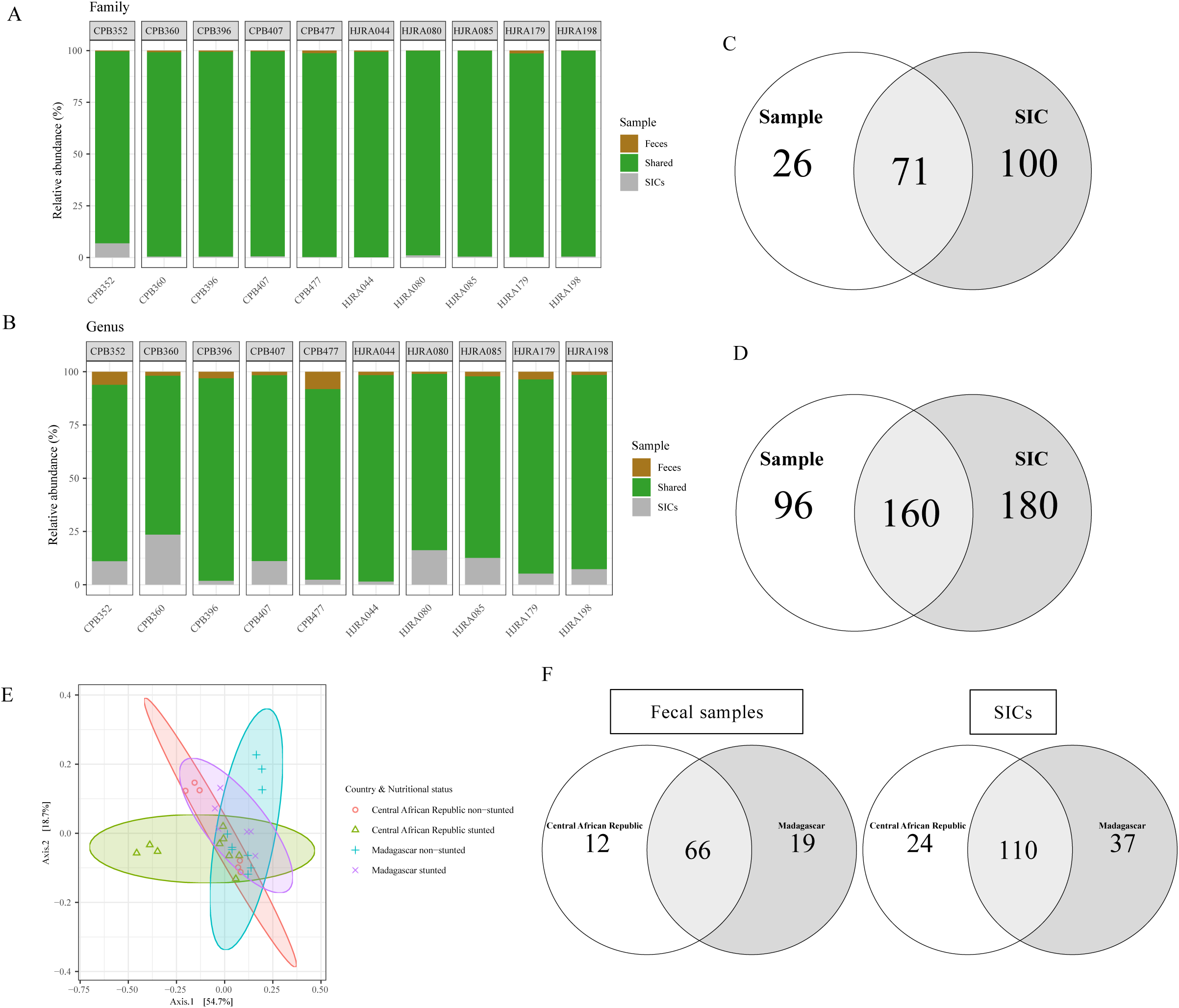
Characterization of the fecal samples and stool-derived *in vitro* communities (SICs). (A) Mean relative abundance of the bacterial families shared between the original fecal samples and their corresponding SIC, unique to the feces, and unique to the SICs. (B) Mean relative abundance of the bacterial genera shared between the original fecal samples and their corresponding SIC, unique to the feces and unique to the SICs. (C) Venn diagram of the bacterial families shared between the fecal samples and SICs and unique to the samples or the SICs. (D) Venn diagram of the bacterial genera shared between the fecal samples and SICs, and unique to the samples or the SICs. (E) PCoA based on Bray-Curtis distance at the family level of the fecal samples. (F) Left: Venn diagram of the bacterial families shared between the fecal samples from Madagascar and Central African Republic and unique to one of the countries. Right: Venn diagram of the bacterial families shared between the SICs from Madagascar and Central African Republic and unique to one of the countries.

**Fig S8:**
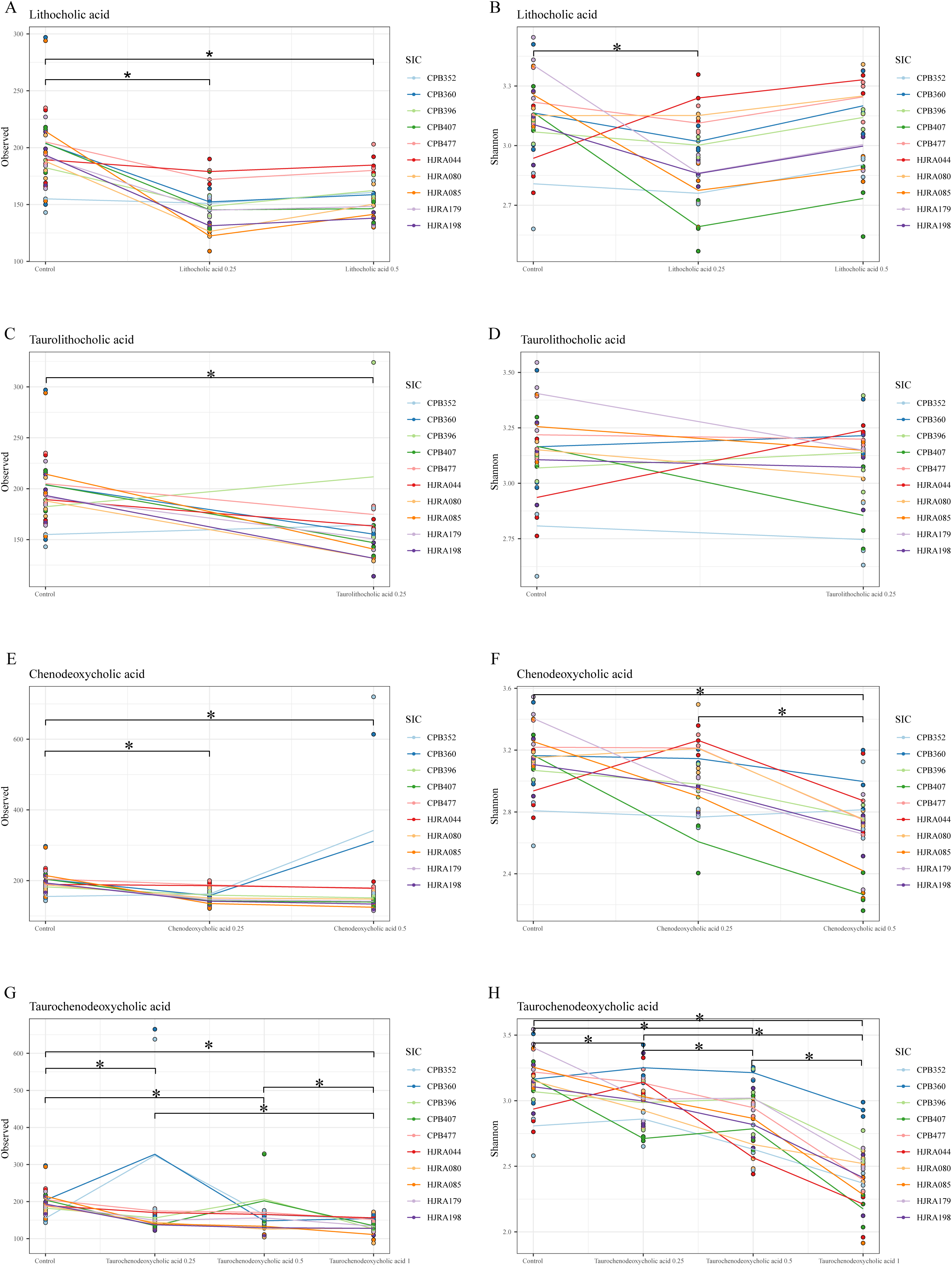
Alpha diversity changes with bile acid treatment (ASV level). (A) Observed index changes of the SICs upon treatment with lithocholic acid. Stars show a significant difference by the Wilcoxon rank-sum test (p<0.05). (B) Shannon index changes of the SICs upon treatment with lithocholic acid. Stars show a significant difference by Wilcoxon rank-sum test (p<0.05). (C) Observed index changes of the SICs upon treatment with taurolithocholic acid. Stars show significant difference by Wilcoxon rank-sum test (p<0.05). (D) Shannon index changes of the SICs upon treatment with taurolithocholic acid. Stars show significant difference by Wilcoxon rank-sum test (p<0.05). (E) Observed index changes of the SICs upon treatment with chenodeoxycholic acid. Stars show significant difference by Wilcoxon rank-sum test (p<0.05). (F) Shannon index changes of the SICs upon treatment with chenodeoxycholic acid. Stars show significant difference by Wilcoxon rank-sum test (p<0.05). (G) Observed index changes of the SICs upon treatment with taurochenodeoxycholic acid. Stars show significant difference by Wilcoxon rank-sum test (p<0.05). (H) Shannon index changes of the SICs upon treatment with taurochenodeoxycholic acid. Stars show significant difference by Wilcoxon rank-sum test (p<0.05).

**Fig S9.**
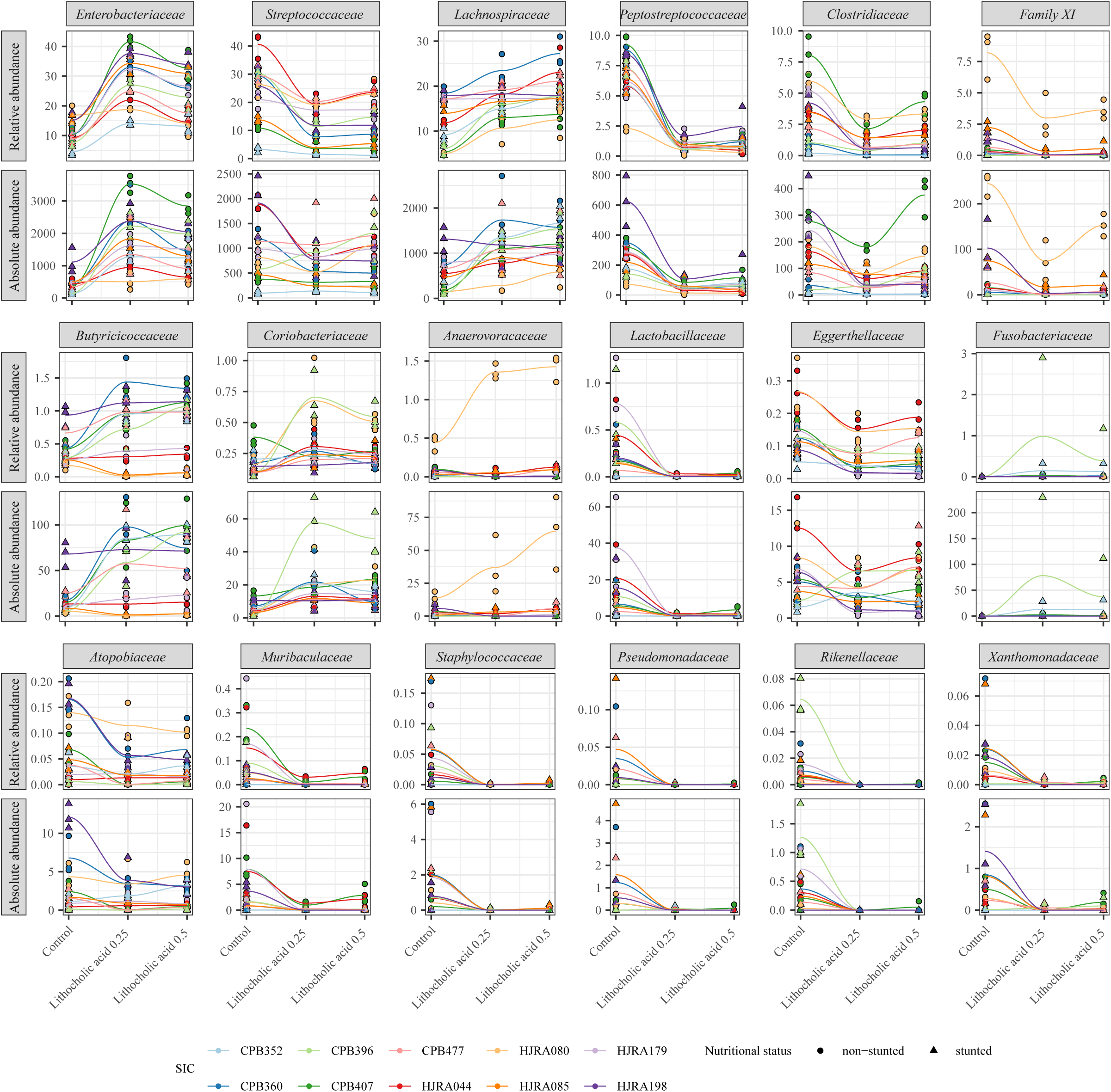
Lithocholic acid boosts butyrate producers in full fecal communities. Change of the relative and absolute abundance of bacterial families identified to be enriched by LefSe, upon treatment with lithocholic acid. Absolute abundance was estimated by multiplying the relative abundance by the DNA concentration of the extracted SICs.

**Fig S10.**
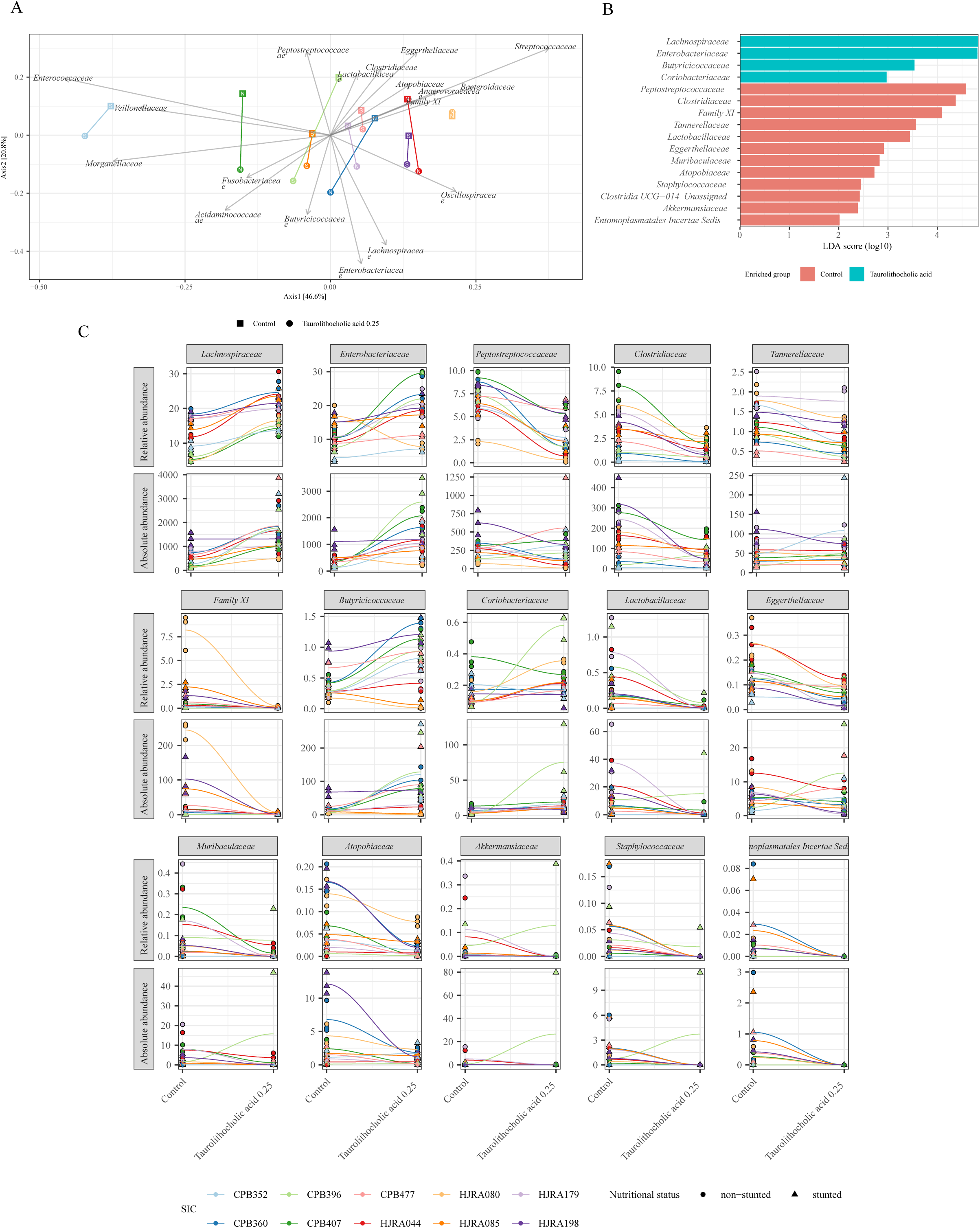
Taurolithocholic acid boosts butyrate producers in full fecal communities. (A) Principal Coordinate Analysis (PCoA) based on Bray-Curtis distance at the family level of the SICs treated with taurolithocholic acid. Each point represents the centroid of the three SIC replicates, and shapes represent the different treatment conditions. SIC from stunted children are depicted with an S inside the shapes. SIC from non-stunted children are depicted with an N inside the shapes. Arrows represent the direction and strength of the significant correlation between bacterial families and the ordination space as identified by envfit (p<0.05). (B) Bacterial families enriched in the control or SIC treated with taurolithocholic acid as identified by LefSe (LDA > 2 and q-value<0.05). (C) Change of the relative and absolute abundance of bacterial families identified to be enriched by LefSe, upon treatment with taurolithocholic acid. Absolute abundance was estimated by multiplying the relative abundance by the DNA concentration of the extracted SICs.

**Fig S11.**
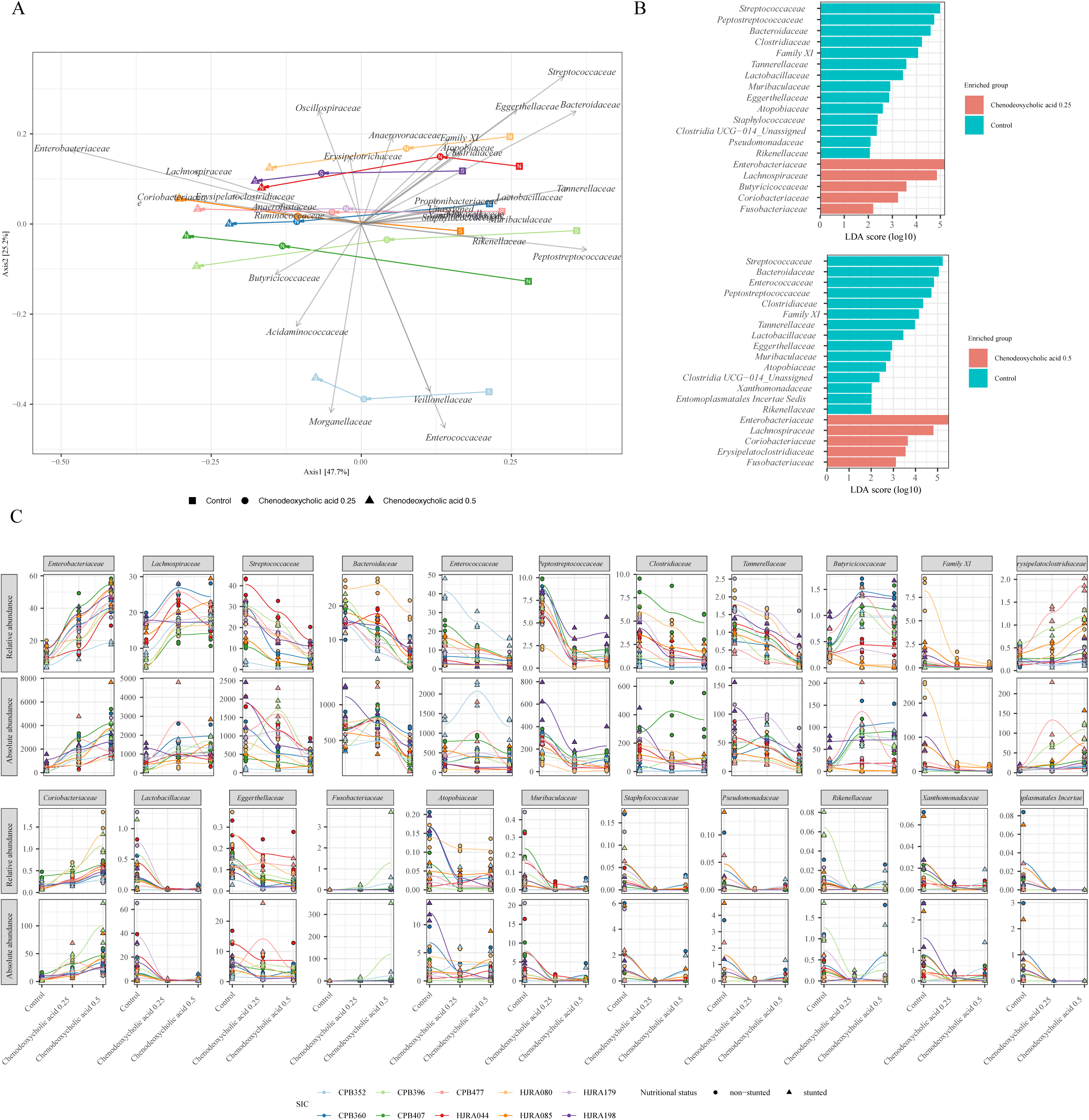
Chenodeoxycholic acid boosts butyrate producers in full fecal communities. (A) Principal Coordinate Analysis (PCoA) based on Bray-Curtis distance at the family level of the SICs treated with chenodeoxycholic acid. Each point represents the centroid of the three SIC replicates, shapes represent the different treatment conditions. SIC from stunted children are depicted with a S inside the shapes. SIC from non-stunted children are depicted with a N inside the shapes. Arrows represent the direction and strength of the significant correlation between bacterial families and the ordination space as identified by envfit (p<0.05). (B) Bacterial families enriched in the control or SIC treated with chenodeoxycholic acid at two different concentrations as identified by LefSe (LDA > 2 and q-value<0.05). (C) Change of the relative and absolute abundance of bacterial families identified to be enriched by LefSe, upon treatment with chenodeoxycholic acid. Absolute abundance was estimated by multiplying the relative abundance by the DNA concentration after extraction.

**Fig S12.**
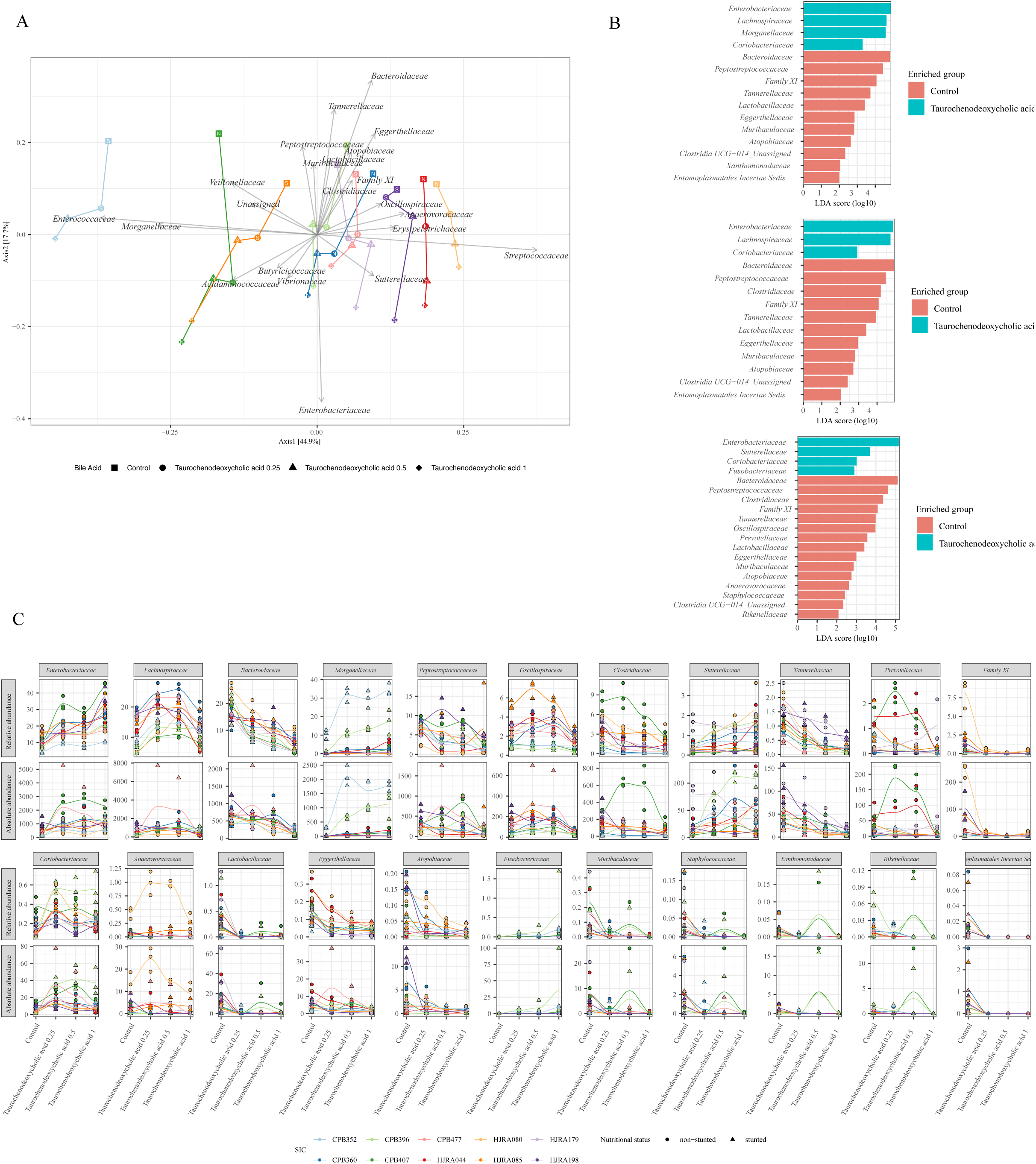
Taurochenodeoxycholic acid boosts butyrate producers in full fecal communities. (A) Principal Coordinate Analysis (PCoA) based on Bray-Curtis distance at the family level of the SICs treated with taurochenodeoxycholic acid. Each point represents the centroid of the three SIC replicates, shapes represent the different treatment conditions. SIC from stunted children are depicted with a S inside the shapes. SIC from non-stunted children are depicted with a N inside the shapes. Arrows represent the direction and strength of the significant correlation between bacterial families and the ordination space as identified by envfit (p<0.05). (B) Bacterial families enriched in the control or SIC treated with taurochenodeoxycholic acid at three different concentrations as identified by LefSe (LDA > 2 and q-value<0.05). (C) Change of the relative and absolute abundance of bacterial families identified to be enriched by LefSe, upon treatment with taurochenodeoxycholic acid. Absolute abundance was estimated by multiplying the relative abundance by the DNA concentration after extraction.

## Supplementary Tables

**Supplementary Table 1: Afribiota 16S rRNA sequencing taxonomy table**

**Supplementary Table 2: Afribiota 16S rRNA sequencing ASV table**

**Supplementary Table 3: Afribiota 16S rRNA sequencing metadata table**

**Supplementary Table 4: Characteristics of reference strains used**

**Supplementary Table 5: Individual Isolate screening AUC results**

**Supplementary Table 6: Synthetic Community screening relative abundance results**

**Supplementary Table 7: Information on stool-derived *in vitro* communities (SICS) and metadata of associated Afribiota donors**

**Supplementary Table 8: Afribiota 16S rRNA sequencing ASV table of fecal samples used for SIC generation**

**Supplementary Table 9: Afribiota 16S rRNA sequencing taxonomy table of fecal samples used for SIC generation**

**Supplementary Table 10: Afribiota 16S rRNA sequencing metadata table of fecal samples used for SIC generation**

**Supplementary Table 11: SIC 16S rRNA sequencing ASV Table**

**Supplementary Table 12: SIC 16S rRNA sequencing taxonomy Table**

**Supplementary Table 13: SIC 16S rRNA sequencing metadata Table**

**Supplementary Table 14: Afribiota dataset: results of non-parametric and linear models on the bile acid levels and cofactors**

**Supplementary Table 15: Clustering results of PCOA drivers**

**Supplementary Table 16: Correlation results of microbiota genera and bile and acids levels in the Afribiota cohort (Spearman)**

**Supplementary Table S17: Chi2 Test Results on Lithochoilc acid and derivatives and microbiota members**

**Supplementary Table S18: Raw data of the growth screening test of individual bacterial taxa**

## List of Afribiota Investigators

The AFRIBIOTA Investigators compose as follows (in alphabetical order):

Jean-Marc Collard, Institut Pasteur de Madagascar, Antananarivo, Madagascar Maria Doria, Institut Pasteur, Paris, France

Darragh Duffy, Institut Pasteur, Paris, France

B. Brett Finlay, University of British Columbia, Vancouver, Canada

Serge Ghislain Djorie, Institut Pasteur de Bangui, Bangui, Central African Republic Tamara Giles-Vernick, Institut Pasteur, Paris, France

Bolmbaye Privat Godje, Complexe Pédiatrique de Bangui, Bangui, Central African Republic

Jean-Chrysostome Gody, Complexe Pédiatrique de Bangui, Bangui, Central African Republic

Milena Hasan, Institut Pasteur, France

Nathalie Kapel, Hôpital Pitié-Salpêtrière, Paris, France

Laurence Trystram-Barbot, Hôpital Pitié-Salpêtrière, Paris, France

Jean-Pierre Lombart, Institut Pasteur de Bangui, Bangui, Central African Republic Alexandre Manirakiza, Institut Pasteur de Bangui, Bangui, Central African Republic Synthia Nazita Nigatoloum, Complexe Pédiatrique de Bangui, Bangui, Central African Republic

Laura Wegener Parfrey, University of British Columbia, Vancouver, Canada Maheninasy Rakotondrainipiana, Institut Pasteur de Madagascar, Antananarivo, Madagascar

Ravaka Niaina Randriamparany, Institut Pasteur de Madagascar, Antananarivo, Madagascar

Prisca Vega Andriantsalama, Institut Pasteur de Madagascar, Antananarivo, Madagascar

Rindra Randremanana, Institut Pasteur de Madagascar, Antananarivo, Madagascar Harifetra Mamy Richard Randriamizao, Centre Hospitalier Universitaire Joseph Pierre-Alain Rubbo, Institut Pasteur de Bangui, Bangui, République Centrafricaine Philippe Sansonetti, Institut Pasteur, Paris, France

Laura Schaeffer, Institut Pasteur, Paris, France

Ionela Gouandjika-Vassilache, Institut Pasteur de Bangui, Bangui, République Centrafricaine

Pascale Vonaesch, Institut Pasteur, Paris, France

Sonia Sandrine Vondo, Complexe Pédiatrique de Bangui, Bangui, Central African Republic

Inès Vigan-Womas, Institut Pasteur de Madagascar, Antananarivo, Madagascar

## Acknowledgments

The authors thank all children and their families involved in the Afribiota project, the participating hospitals in Bangui and Antananarivo, Institut Pasteur, Institut Pasteur de Madagascar, and Institut Pasteur de Bangui and members of the scientific advisory board for their continuous support for the Afribiota project. Further, they thank the Centre de Recherche Translationelle and the Direction Internationale of the Institut Pasteur, and especially Paméla Palvadeau, Jane Lynda Deuve, Cécile Artaud, Nathalie Jolly, Sophie Jarrijon, Mamy Ratsialonina, and Jean-François Damaras for their precious help in setting up and steering the AFRIBIOTA project. The authors also acknowledge Jean-Marc Collard, Pierre-Alain Rubbo, Dieu-Merci Welekoi-Yapondo, Lova Andrianonimiadana, Laurence Arowas and Marie-Noelle Ungeheuer for managing the AFRIBIOTA biobank. We last thank Rizlan Bernier-Latmani for helpful discussions and for sharing their grouping of the bile acids.

## References

1 Mohanty, I. et al. The changing metabolic landscape of bile acids - keys to metabolism and immune regulation. Nat Rev Gastroenterol Hepatol 21, 493–516, doi:10.1038/s41575-024-00914-3 (2024).

2 Agus, A., Clement, K. & Sokol, H. Gut microbiota-derived metabolites as central regulators in metabolic disorders. Gut 70, 1174–1182, doi:10.1136/gutjnl-2020-323071 (2021).

3 Monteiro-Cardoso, V. F., Corlianò, M. & Singaraja, R. R. Bile Acids: A Communication Channel in the Gut-Brain Axis. Neuromol Med 23, 99–117, doi:10.1007/s12017-020-08625-z (2021).

4 Wahlström, A., Sayin, S. I., Marschall, H.-U. & Bäckhed, F. Intestinal Crosstalk between Bile Acids and Microbiota and Its Impact on Host Metabolism. Cell metabolism 24, 41–50, doi:10.1016/j.cmet.2016.05.005 (2016).

5 Dahl, W. J., Rivero Mendoza, D. & Lambert, J. M. Diet, nutrients and the microbiome. Prog Mol Biol Transl Sci 171, 237–263, doi:10.1016/bs.pmbts.2020.04.006 (2020).

6 Joyce, S. A. et al. Regulation of host weight gain and lipid metabolism by bacterial bile acid modification in the gut. P Natl Acad Sci USA 111, 7421–7426, doi:10.1073/pnas.1323599111 (2014).

7 Wang, K. et al. Parabacteroides distasonis Alleviates Obesity and Metabolic Dysfunctions via Production of Succinate and Secondary Bile Acids. Cell reports 26, 222–235.e225, doi:10.1016/j.celrep.2018.12.028 (2019).

8 Natividad, J. M. et al. Bilophila wadsworthia aggravates high fat diet induced metabolic dysfunctions in mice. Nature communications 9, 2802, doi:10.1038/s41467-018-05249-7 (2018).

9 Hasan, F. et al. Bile acid dysmetabolism in Bangladeshi infants is associated with poor linear growth, enteric inflammation, and small intestine bacterial overgrowth. medRxiv, doi:10.1101/2025.02.04.25321650 (2025).

10 Zhang, L. et al. Impaired Bile Acid Homeostasis in Children with Severe Acute Malnutrition. PLoS One 11, e0155143, doi:10.1371/journal.pone.0155143 (2016).

11 Watanabe, M., Fukiya, S. & Yokota, A. Comprehensive evaluation of the bactericidal activities of free bile acids in the large intestine of humans and rodents. Journal of lipid research 58, 1143–1152, doi:10.1194/jlr.M075143 (2017).

12 Larabi, A. B., Masson, H. L. P. & Baumler, A. J. Bile acids as modulators of gut microbiota composition and function. Gut Microbes 15, 2172671, doi:10.1080/19490976.2023.2172671 (2023).

13 Parks, R. W. et al. Intestinal barrier dysfunction in clinical and experimental obstructive jaundice and its reversal by internal biliary drainage. Brit J Surg 83, 1345–1349, doi:DOI 10.1002/bjs.1800831007 (1996).

14 Lorenzo-Zúñiga, V. et al. Oral bile acids reduce bacterial overgrowth, bacterial translocation, and endotoxemia in cirrhotic rats. Hepatology (Baltimore, Md.) 37, 551–557, doi:10.1053/jhep.2003.50116 (2003).

15 Eade, C. R. et al. Bile Acids Function Synergistically To Repress Invasion Gene Expression in Salmonella by Destabilizing the Invasion Regulator HilD. Infection and immunity 84, 2198–2208, doi:10.1128/IAI.00177-16 (2016).

16 Bachmann, V. et al. Bile Salts Modulate the Mucin-Activated Type VI Secretion System of Pandemic Vibrio cholerae. PLoS neglected tropical diseases 9, e0004031, doi:10.1371/journal.pntd.0004031 (2015).

17 Yang, M. et al. Bile salt-induced intermolecular disulfide bond formation activates Vibrio cholerae virulence. P Natl Acad Sci USA 110, 2348–2353, doi:10.1073/pnas.1218039110 (2013).

18 Antunes, L. C. M. et al. Repression of Salmonella enterica phoP expression by small molecules from physiological bile. Journal of bacteriology 194, 2286–2296, doi:10.1128/JB.00104-12 (2012).

19 Studer, N. et al. Functional Intestinal Bile Acid 7α-Dehydroxylation by Clostridium scindens Associated with Protection from Clostridium difficile Infection in a Gnotobiotic Mouse Model. Frontiers in cellular and infection microbiology 6, 191, doi:10.3389/fcimb.2016.00191 (2016).

20 Thanissery, R., Winston, J. A. & Theriot, C. M. Inhibition of spore germination, growth, and toxin activity of clinically relevant C. difficile strains by gut microbiota derived secondary bile acids. Anaerobe 45, 86–100, doi:10.1016/j.anaerobe.2017.03.004 (2017).

21 Theriot, C. M., Bowman, A. A. & Young, V. B. Antibiotic-Induced Alterations of the Gut Microbiota Alter Secondary Bile Acid Production and Allow for Clostridium difficile Spore Germination and Outgrowth in the Large Intestine. mSphere 1, e00045–00015, doi:10.1128/mSphere.00045-15 (2016).

22 Winston, J. A. & Theriot, C. M. Impact of microbial derived secondary bile acids on colonization resistance against Clostridium difficile in the gastrointestinal tract. Anaerobe 41, 44–50, doi:10.1016/j.anaerobe.2016.05.003 (2016).

23 Duboc, H. et al. Connecting dysbiosis, bile-acid dysmetabolism and gut inflammation in inflammatory bowel diseases. Gut 62, 531–539, doi:10.1136/gutjnl-2012-302578 (2013).

24 Duboc, H. et al. Increase in fecal primary bile acids and dysbiosis in patients with diarrhea-predominant irritable bowel syndrome. Neurogastroent Motil 24, 513-+, doi:10.1111/j.1365-2982.2012.01893.x (2012).

25 Van den Bossche L., et al. Ursodeoxycholic Acid and Its Taurine- or Glycine-Conjugated Species Reduce Colitogenic Dysbiosis and Equally Suppress Experimental Colitis in Mice. Applied and environmental microbiology 83, e02766–02716, doi:10.1128/AEM.02766-16 (2017).

26 Lloyd-Price, J. et al. Multi-omics of the gut microbial ecosystem in inflammatory bowel diseases. Nature 569, 655–662, doi:10.1038/s41586-019-1237-9 (2019).

27. Local Burden of Disease Child Growth Failure, C. Mapping child growth failure across low- and middle-income countries. Nature 577, 231–234, doi:10.1038/s41586-019-1878-8 (2020).

28 Chibuye, M. et al. Systematic review of associations between gut microbiome composition and stunting in under-five children. Npj Biofilms Microbi 10, doi:ARTN 46 10.1038/s41522-024-00517-5 (2024).

29 Chen, R. Y. et al. Duodenal Microbiota in Stunted Undernourished Children with Enteropathy. N Engl J Med 383, 321–333, doi:10.1056/NEJMoa1916004 (2020).

30 Collard, J. M. et al. High prevalence of small intestine bacteria overgrowth and asymptomatic carriage of enteric pathogens in stunted children in Antananarivo, Madagascar. PLoS Negl Trop Dis 16, e0009849, doi:10.1371/journal.pntd.0009849 (2022).

31 Vonaesch, P. et al. Stunted children display ectopic small intestinal colonization by oral bacteria, which cause lipid malabsorption in experimental models. Proc Natl Acad Sci U S A 119, e2209589119, doi:10.1073/pnas.2209589119 (2022).

32 Vonaesch, P. et al. Stunted childhood growth is associated with decompartmentalization of the gastrointestinal tract and overgrowth of oropharyngeal taxa. P Natl Acad Sci USA 115, E8489–E8498, doi:10.1073/pnas.1806573115 (2018).

33 Gough, E. K. et al. Linear growth faltering in infants is associated with Acidaminococcus sp. and community-level changes in the gut microbiota. Microbiome 3, 24, doi:10.1186/s40168-015-0089-2 (2015).

34 Bagamian, K. H. et al. Could a Shigella vaccine impact long-term health outcomes?: Summary report of an expert meeting to inform a Shigella vaccine public health value proposition, March 24 and 29, 2021. Vaccine X 12, 100218, doi:10.1016/j.jvacx.2022.100218 (2022).

35 Crane, R. J., Jones, K. D. & Berkley, J. A. Environmental enteric dysfunction: an overview. Food Nutr Bull 36, S76–87, doi:10.1177/15648265150361S113 (2015).

36 Vonaesch, P. et al. Putative Biomarkers of Environmental Enteric Disease Fail to Correlate in a Cross-Sectional Study in Two Study Sites in Sub-Saharan Africa. Nutrients 14, doi:10.3390/nu14163312 (2022).

37 Haberman, Y. et al. Mucosal Genomics Implicate Lymphocyte Activation and Lipid Metabolism in Refractory Environmental Enteric Dysfunction. Gastroenterology 160, 2055–2071 e2050, doi:10.1053/j.gastro.2021.01.221 (2021).

38 Harper, K. M., Mutasa, M., Prendergast, A. J., Humphrey, J. & Manges, A. R. Environmental enteric dysfunction pathways and child stunting: A systematic review. PLoS neglected tropical diseases 12, e0006205, doi:10.1371/journal.pntd.0006205 (2018).

39 Semba, R. D. et al. Environmental Enteric Dysfunction Is Associated With Altered Bile Acid Metabolism. Journal of pediatric gastroenterology and nutrition 64, 536–540, doi:10.1097/MPG.0000000000001313 (2017).

40 Zhao, X. et al. Bile Acid Profiling Reveals Distinct Signatures in Undernourished Children with Environmental Enteric Dysfunction. J Nutr 151, 3689–3700, doi:10.1093/jn/nxab321 (2021).

41 Moreau, G. B. et al. Childhood growth and neurocognition are associated with distinct sets of metabolites. EBioMedicine 44, 597–606, doi:10.1016/j.ebiom.2019.05.043 (2019).

42 Vonaesch, P. et al. Identifying the etiology and pathophysiology underlying stunting and environmental enteropathy: study protocol of the AFRIBIOTA project. BMC pediatrics 18, 236, doi:10.1186/s12887-018-1189-5 (2018).

43 Vonaesch, P. et al. Factors Associated with Stunted Growth in Children Under Five Years in Antananarivo, Madagascar and Bangui, Central African Republic. Matern Child Health J 25, 1626–1637, doi:10.1007/s10995-021-03201-8 (2021).

44 Razanajatovo, I. M. et al. Factors Associated with Carriage of Enteropathogenic and Non-Enteropathogenic Viruses: A Reanalysis of Matched Case-Control Data from the AFRIBIOTA Site in Antananarivo, Madagascar. Pathogens 12, doi:10.3390/pathogens12081009 (2023).

45 Habib, A. et al. High prevalence of intestinal parasite infestations among stunted and control children aged 2 to 5 years old in two neighborhoods of Antananarivo, Madagascar. PLoS Negl Trop Dis 15, e0009333, doi:10.1371/journal.pntd.0009333 (2021).

46 Huus, K. E. et al. Immunoglobulin recognition of fecal bacteria in stunted and non-stunted children: findings from the Afribiota study. Microbiome 8, 113, doi:10.1186/s40168-020-00890-1 (2020).

47 Vonaesch, P. et al. The eukaryome of African children is influenced by geographic location, gut biogeography, and nutritional status. Microlife 4, uqad033, doi:10.1093/femsml/uqad033 (2023).

48. Group, W. M. G. R. S. … Standards: Head circumference-for-age, arm circumference-for-age, triceps skinfold-for-age and subscapular skinfold-for-age: Methods and development. (Geneva: World Health …, 2007).

49 Kozich, J. J., Westcott, S. L., Baxter, N. T., Highlander, S. K. & Schloss, P. D. Development of a dual-index sequencing strategy and curation pipeline for analyzing amplicon sequence data on the MiSeq Illumina sequencing platform. Applied and environmental microbiology 79, 5112–5120, doi:10.1128/AEM.01043-13 (2013).

50 Callahan, B. J. et al. DADA2: High-resolution sample inference from Illumina amplicon data. Nat Methods 13, 581–583, doi:10.1038/nmeth.3869 (2016).

51 Hartman, S. J. et al. A microbiome-directed therapeutic food for children recovering from severe acute malnutrition. medRxiv, doi:10.1101/2024.06.11.24307076 (2024).

52 Maier, L. et al. Extensive impact of non-antibiotic drugs on human gut bacteria. Nature 555, 623–628, doi:10.1038/nature25979 (2018).

53 Wright, E. S. Using DECIPHER v2.0 to Analyze Big Biological Sequence Data in R. R J 8, 352–359 (2016).

54 Leek, J. T., Johnson, W. E., Parker, H. S., Jaffe, A. E. & Storey, J. D. The sva package for removing batch effects and other unwanted variation in high-throughput experiments. Bioinformatics 28, 882–883, doi:10.1093/bioinformatics/bts034 (2012).

55 McMurdie, P. J. & Holmes, S. phyloseq: an R package for reproducible interactive analysis and graphics of microbiome census data. PloS one 8, e61217, doi:10.1371/journal.pone.0061217 (2013).

56 Callahan, B. J., Sankaran, K., Fukuyama, J. A., McMurdie, P. J. & Holmes, S. P. Bioconductor Workflow for Microbiome Data Analysis: from raw reads to community analyses. F1000Res 5, 1492, doi:10.12688/f1000research.8986.2 (2016).

57 Liu, C., Cui, Y., Li, X. & Yao, M. microeco: an R package for data mining in microbial community ecology. FEMS Microbiol Ecol 97, doi:10.1093/femsec/fiaa255 (2021).

58 Dixon, P. VEGAN, a package of R functions for community ecology. Journal of Vegetation Science 14, 927–930, doi:10.1111/j.1654-1103.2003.tb02228.x (2009).

59 Paradis, E., Claude, J. & Strimmer, K. APE: Analyses of Phylogenetics and Evolution in R language. Bioinformatics 20, 289–290, doi:10.1093/bioinformatics/btg412 (2004).

60 Softw, H. W. J. S. & 2010. ggplot2: elegant graphics for data analysis. researchgate.net.

61 Cao, Y. et al. microbiomeMarker: an R/Bioconductor package for microbiome marker identification and visualization. Bioinformatics 38, 4027–4029, doi:10.1093/bioinformatics/btac438 (2022).

62 Sistrunk, J. R., Nickerson, K. P., Chanin, R. B., Rasko, D. A. & Faherty, C. S. Survival of the Fittest: How Bacterial Pathogens Utilize Bile To Enhance Infection. Clin Microbiol Rev 29, 819–836, doi:10.1128/CMR.00031-16 (2016).

63. 63 Garneau, J. R. et al. Phage Endolysin Enables Targeted Manipulation of the Small Intestinal Microbiota and Uncovers Niche Overlap Between Oral and Butyrate-Producing Taxa. bioRxiv, 2026.2001.2006.697609, doi:10.64898/2026.01.06.697609 (2026).

64 Preidis, G. A. et al. The undernourished neonatal mouse metabolome reveals evidence of liver and biliary dysfunction, inflammation, and oxidative stress. J Nutr 144, 273–281, doi:10.3945/jn.113.183731 (2014).

65 Brown, E. M. et al. Diet and specific microbial exposure trigger features of environmental enteropathy in a novel murine model. Nature communications 6, 7806, doi:10.1038/ncomms8806 (2015).

66 van Best, N. et al. Bile acids drive the newborn’s gut microbiota maturation. Nature Communications 11, doi:ARTN 3692 10.1038/s41467-020-17183-8 (2020).

67 Subramanian, S. et al. Persistent gut microbiota immaturity in malnourished Bangladeshi children. Nature 510, 417–421, doi:10.1038/nature13421 (2014).

68 Heinken, A. et al. Systematic assessment of secondary bile acid metabolism in gut microbes reveals distinct metabolic capabilities in inflammatory bowel disease. Microbiome 7, 75, doi:10.1186/s40168-019-0689-3 (2019).

69 Sinha, S. R. et al. Dysbiosis-Induced Secondary Bile Acid Deficiency Promotes Intestinal Inflammation. Cell Host Microbe 27, 659–670 e655, doi:10.1016/j.chom.2020.01.021 (2020).

70 Qu, Q. et al. Lithocholic acid phenocopies anti-ageing effects of calorie restriction. Nature, doi:10.1038/s41586-024-08329-5 (2024).

71 Alavi, S. et al. Interpersonal Gut Microbiome Variation Drives Susceptibility and Resistance to Cholera Infection. Cell 181, 1533–1546 e1513, doi:10.1016/j.cell.2020.05.036 (2020).

72 Sutherland, J. D. & Macdonald, I. A. The metabolism of primary, 7-oxo, and 7 beta-hydroxy bile acids by Clostridium absonum. J Lipid Res 23, 726–732 (1982).

73 Lepercq, P. et al. Increasing ursodeoxycholic acid in the enterohepatic circulation of pigs through the administration of living bacteria. Br J Nutr 93, 457–469, doi:10.1079/bjn20041386 (2005).

74 Lee, J. Y. et al. Contribution of the 7beta-hydroxysteroid dehydrogenase from Ruminococcus gnavus N53 to ursodeoxycholic acid formation in the human colon. J Lipid Res 54, 3062–3069, doi:10.1194/jlr.M039834 (2013).

75 Alnouti, Y. Bile Acid sulfation: a pathway of bile acid elimination and detoxification. Toxicol Sci 108, 225–246, doi:10.1093/toxsci/kfn268 (2009).

76 Kriaa, A. et al. Microbial impact on cholesterol and bile acid metabolism: current status and future prospects. J Lipid Res 60, 323–332, doi:10.1194/jlr.R088989 (2019).

77 Peng, Y. L. et al. Effects of bile acids on the growth, composition and metabolism of gut bacteria. NPJ Biofilms Microbiomes 10, 112, doi:10.1038/s41522-024-00566-w (2024).

78 Friedman, E. S. et al. FXR-Dependent Modulation of the Human Small Intestinal Microbiome by the Bile Acid Derivative Obeticholic Acid. Gastroenterology 155, 1741–1752 e1745, doi:10.1053/j.gastro.2018.08.022 (2018).

79 Kurdi, P., Kawanishi, K., Mizutani, K. & Yokota, A. Mechanism of growth inhibition by free bile acids in lactobacilli and bifidobacteria. J Bacteriol 188, 1979–1986, doi:10.1128/JB.188.5.1979-1986.2006 (2006).

80 Nickerson, K. P. et al. Analysis of Shigella flexneri Resistance, Biofilm Formation, and Transcriptional Profile in Response to Bile Salts. Infect Immun 85, doi:10.1128/IAI.01067-16 (2017).

